# CHCHD4 confers metabolic vulnerabilities to tumour cells through its control of the mitochondrial respiratory chain

**DOI:** 10.1101/513770

**Authors:** Luke W. Thomas, Jenna M. Stephen, Cinzia Esposito, Simon Hoer, Robin Antrobus, Afshan Ahmed, Hasan Al-Habib, Margaret Ashcroft

**Affiliations:** Department of Medicine, University of Cambridge, Cambridge Biomedical Campus, Cambridge, CB2 0AH. United Kingdom; Cambridge Institute for Medical Research, University of Cambridge, Cambridge Biomedical Campus, Cambridge, CB2 0XY. United Kingdom

**Keywords:** Coiled-coil-helix-coiled-coil-helix domain containing 4 (CHCHD4), hypoxia, HIF-1α, mitochondria, respiratory chain, disulfide relay system, complex I, tumour growth, tumour metabolism

## Abstract

**BACKGROUND:** Tumour cells rely on glycolysis and mitochondrial oxidative phosphorylation (OXPHOS) to survive. Thus mitochondrial OXPHOS has become an increasingly attractive area for therapeutic exploitation in cancer. However, mitochondria are required for intracellular oxygenation and normal physiological processes, and it remains unclear which mitochondrial molecular mechanisms might provide therapeutic benefit. Previously, we discovered that coiled-coil helix coiled-coil helix domain-containing protein 4 (CHCHD4) is critical for maintaining intracellular oxygenation and required for the cellular response to hypoxia (low oxygenation) in tumour cells through molecular mechanisms that we do not yet fully understand. Overexpression of *CHCHD4* in human cancers, correlates with increased tumour progression and poor patient survival.

**RESULTS:** Here, we show that elevated CHCHD4 expression provides a proliferative and metabolic advantage to tumour cells in normoxia and hypoxia. Using stable isotope labelling with amino acids in cell culture (SILAC) and analysis of the whole mitochondrial proteome, we show that CHCHD4 dynamically affects the expression of a broad range of mitochondrial respiratory chain subunits from complex I-V, including multiple subunits of complex I (CI) required for complex assembly that are essential for cell survival. We found that loss of CHCHD4 protects tumour cells from respiratory chain inhibition at CI, while elevated CHCHD4 expression in tumour cells leads to significantly increased sensitivity to CI inhibition, in part through the production of mitochondrial reactive oxygen species (ROS).

**CONCLUSIONS:** Our study highlights an important role for CHCHD4 in regulating tumour cell metabolism, and reveals that CHCHD4 confers metabolic vulnerabilities to tumour cells through its control of the mitochondrial respiratory chain and CI biology.

## BACKGROUND

Metabolic reprogramming and altered mitochondrial metabolism is a feature of cancer, and thus has become an attractive area for therapeutic exploitation [1, 2]. Given the importance of mitochondria in controlling normal physiological processes, understanding how mitochondrial metabolism underlies tumorigenesis is important for ascertaining its therapeutic potential in cancer.

Previously, we discovered the essential redox-sensitive mitochondrial intermembrane space (IMS) protein CHCHD4, is a critical for regulating intracellular oxygen consumption rate and metabolic responses to low oxygen (hypoxia) in tumour cells [3, 4]. Overexpression of *CHCHD4* in human cancers significantly correlates with the hypoxia gene signature, tumour progression, disease recurrence and poor patient survival [3].

CHCHD4 provides an import and oxidoreductase-mediated protein folding function along with the sulfhydryl oxidase GFER (ALR/Erv1) as a key part of the disulphide relay system (DRS) within the mitochondrial IMS [5–7]. As such, CHCHD4 controls the import of a number of mitochondrial proteins that contain a twin-CX_9_C or twin-CX_3_C motif [8–10]. Additionally, as a component of the DRS, CHCHD4 participates in electron transfer to CIV, the molecular oxygen acceptor of the respiratory chain [11]. We and others have found that the functionally conserved cysteines within the redox sensitive Cys-Pro-Cys (CPC) domain of CHCHD4 regulate its mitochondrial localisation in yeast [12–14] and human cells [3, 15]. Recently, we discovered that CHCHD4 regulates intracellular oxygenation in tumour cells, which is dependent on the functionally important cysteines of the CPC motif and CIV activity [4].

In this study, using both loss- and gain-of-function approaches, we have further explored the mitochondrial mechanism(s) by which CHCHD4 regulates respiratory chain function and tumour cell metabolism.

## METHODS

### Cell culture

Human osteosarcoma U2OS control and independent clonal cell lines (WT.cl1 and WT.cl3) expressing CHCHD4.1 cDNA (CHCHD4-WT-expressing cells) or CHCHD4-C66A/C668A cDNA (CHCHD4-(C66A/C68A)-expressing cells) have been described by us recently [4]. Human U2OS-HRE-luc [16] or human HCT116 colon carcinoma cells [17] were used to stably express two independent shRNA control vectors (Empty Vector (shRNA control 1) and GFP Vector (shRNA control 2)) or two independent shRNAs targeting CHCHD4 (CHCHD4 shRNA1 or CHCHD4 shRNA2) utilizing a green fluorescent protein (GFP)-SMARTvector™ pre-packaged lentivirus system from ThermoFisher Scientific. Independent cell lines were selected, expanded and characterized. All cell lines were maintained in Dulbecco’s modified eagle medium (DMEM) containing 4.5g/L glucose (#41966-029, Life Technologies), and supplemented with 10% fetal calf serum (#EU-000-F, SeraLabs), 100 IU/mL penicillin/100 μg/mL streptomycin (#15140-122, Life Technologies) and 6 mM L-glutamine (#25030-024, Life Technologies). Cell lines used were authenticated and routinely confirmed to be negative for any mycoplasma contamination. Hypoxia was achieved by incubating cells in 1% O_2_, 5% CO_2_ and 94% N_2_ in a Ruskinn SCI-tive workstation, without agitation.

### Antibodies and reagents

For antibodies, the catalogue number and working dilution used are indicated in brackets. The rabbit polyclonal CHCHD4 (HPA34688, 1:1000) antibody was purchased from Cambridge Biosciences. The mouse monoclonal HIF-1α antibody (#610959, 1:500) was purchased from BD Biosciences. The mouse monoclonal β-actin (ab6276, 1:10000), mouse monoclonal α-Tubulin (ab7291, 1:1000), rabbit polyclonal NDUFS3 (ab110246, 1:500) and rabbit polyclonal UQCRC2 (ab14745, 1:1000) were purchased from Abcam. The mouse monoclonal anti-myc (9B11 clone, #2276, 1:1000)), rabbit polyclonal PHB1 (#2426, 1:1000), rabbit polyclonal HIF-2α (#7096, 1:1000), rabbit polyclonal AIF (#4642, 1:1000), rabbit polyclonal SDHA (#11998, 1:1000) and rabbit monoclonal COXIV (3E11 clone, #4850, 1:1000) antibodies were purchased from Cell Signaling Technology. The donkey anti-rabbit (NA934, 1:1000) and anti-mouse (NA931, 1:1000) horseradish peroxidase (HRP)-linked secondary antibodies were purchased from VWR. The GFER rabbit polyclonal antibody (#HPA041227, 1:1000), oligomycin (#75351), rotenone (#R8875), FCCP (#C2920), TMRM (#T5428), D-glucose (#G8270), 2-deoxy-D-glucose (#D8375) and sodium azide (NaAzide, #S8032) were purchased from Sigma Aldrich. MitoSOX Red (#M36008) was purchased from ThermoFisher Scientific. MitoView Green (#70054) was purchased from Biotium. 2-NBDG (#N13195) was purchased from Life Technologies. Trolox (#648471) was purchased from Merck Millipore. BAY 87-2243 (#HY-15836) was purchased from MedChemTronica AB. NSC-134754, 3-Ethyl-9,10-dimethoxy-2-(1,2,3,4-tetrahydro-isoquinolin-1-ylmethyl)-1,6,7,11b-tetrahydro-4H-pyrido[2,1-a]isoquinoline and NSC-134756 were obtained from the National Cancer Centre, Drug Therapeutic Program (NCI-DTP), Frederick MD and dissolved in dimethyl sulfoxide (DMSO) as we have described previously [16, 18–20].

### Gene silencing

Non-silencing siRNA duplexes (MISSION^®^ siRNA Universal Negative Control #1, SIC-001) and custom designed siRNA duplexes were purchased from Sigma-Aldrich, and transfected into subconfluent cells using HiPerfect transfection (QIAGEN) according to the manufacturer’s instructions. The target sequences for CHCHD4 were 5’-GAGGAAACGTTGTGAATTA-3’ (siRNA1) and 5’-AAGATTTGGACCCTTCCATTC-3’ (siRNA2). The target sequences for HIF1A were 5’-TACGTTGTGAGTGGTATTATT-3’ (siRNA1) and 5’-TAGAAGGTATGTGGCATTTAT-3’ (siRNA2) (Additional file 7). For the generation of stable shRNA expressing cells, adherent HCT116, or U2OS-HRE-luc cells were incubated in DMEM containing 50 mg/mL Hexadimethrine bromide (polybrene, #H9268, Sigma-Aldrich). Immediately following addition of polybrene-containing DMEM, lentiviral particles were added containing either a non-shRNA expressing vector (shRNA control 1), a non-shRNA GFP expressing vector (shRNA control 2), or vectors expressing CHCHD4 targeted shRNA sequences (CHCHD4 shRNA1, CHCHD4 shRNA2). All lentiviral particles purchased from Dharmacon (ThermoFisher Scientific) and target sequences are detailed in Additional file 7. After 72h of incubation with lentiviral particles, media replaced with maintenance DMEM (w/o polybrene/lentivirus), and cells incubated for a further 72h. Selection of transduced cell pools carried out by the addition of 0.5 μg/mL puromycin (#P9620, Sigma- Aldrich) to the culture medium, followed by incubation for >96h. Knock-down confirmed by both western blotting for CHCHD4 protein expression and QPCR for *CHCHD4* transcript expression.

### Gene expression analysis

Total RNA samples were isolated using the GeneElute kit (#RTN350), following the manufacturer’s protocol (Sigma-Aldrich). cDNA synthesis was carried out using the qScript synthesis kit (#95048-100), following the manufacturer’s protocol (Quantabio). mRNA expression was measured by quantitative (Q)-PCR using SYBR Green Mastermix (#RT-SY2X-NRWOU+B, Eurogentec Ltd.) and the DNA Engine Opticon 2 system (BioRad). The Q-PCR primer sequences are included in Additional file 8.

### Mitochondrial copy number

Measurement of mitochondrial (mt)DNA copy number has been described previously [21]. Total DNA samples were isolated using the QIAamp DNA Blood Mini Kit, following the manufacturer’s protocol (Sigma-Aldrich). Relative copy numbers of the single-copy nuclear-encoded gene beta-2-microglobulin (β2M) and the mitochondrially-encoded gene mtND1 were measured by quantitative (Q)-PCR using SYBR Green Mastermix (Eurogentec Ltd.) and the DNA Engine Opticon 2 system (BioRad). The Q-PCR primer sequences are in Additional file 8. To determine the mtDNA content, relative to nuclear DNA, the following equations were used:

a. ΔC_T_ = (nucDNA C_T_ − mtDNA C_T_)
b. Relative mtDNA content = 2 × 2^ΔCT^

### Respirometry

Oxygen consumption rates (OCR) and extracellular acidification rates (ECAR) were determined using a Seahorse XF96 Analyser (Seahorse Bioscience). Respiratory profiles were generated by serial treatment with optimised concentrations of oligomycin (1 μg/mL), p-[trifluoromethoxy]-phenyl-hydrazone (FCCP, 500 nM), and rotenone (500 nM). Glycolytic profiles were generated by serial treatment of glucose-restricted cells with optimised concentrations of glucose (12.5 mM), oligomycin A (1 μM,) and 2-DG (50 mM). Cell number normalisation was carried out post-respirometry using sulforhodamine B (SRB) staining of TCA fixed cells in the assay plate.

### Mitochondrial fractionation

Crude mitochondrial fractions were prepared from cultured cells as follows. All tubes and reagents were pre-chilled, and all steps carried out at 4°C, or on ice. Cells were collected, and washed twice with homogenisation buffer (HB) (250 mM Mannitol, 5 mM HEPES (pH 7.4), 0.5 mM EGTA, in water). Pellets were resuspended in 1 mL of HB, and transferred to a chilled glass potter. Cells were lysed with 150 strokes of potter on ice, and 50 μL of homogenate removed for whole cell lysate (WCL) sample. The remaining lysate was spun at 1,000 g for 5 min at 4°C. Supernatants were transferred to fresh tubes, and spun at 2,000 g for 5 min at 4°C. Supernatants were again transferred to fresh tubes, and spun at 10,000 g for 10 min at 4°C. 50 μL supernatants were retained as the cytoplasm sample. Mitochondrial pellets were washed with 2-3 mL HB, and spun at 10,000 g for 10 min at 4°C. Supernatants were carefully removed, and mitochondrial pellets were resuspended in 200 μL HB for functional assays, or 200 μL 1x Laemmli sample buffer for immunoblotting. The functionality of isolated mitochondria was monitored using TMRM staining, which measures mitochondrial membrane potential (Additional file 2g).

### SILAC

Cells were incubated in arginine and lysine free DMEM (#A14431, Life Technologies), supplemented with either (light) L-lysine (#L8662) and L-arginine (#A5006), or (heavy) L-lysine-^13^C_6_, ^15^N_2_ (Lys-8, #608041)) and L-arginine-^13^C_6_, ^15^N_4_ (Arg-10, #608033) stable isotope labelled amino acids (Sigma-Aldrich). Media was also supplemented with 10% dialysed FCS (Sigma), penicillin (100 IU/mL), streptomycin (100 μg/mL) and L-glutamine (200 mM), all purchased from Life Technologies. Amino acid incorporation was carried out over >5 passages. Mitochondrial fractions were isolated as described below. 50 μg of enriched mitochondria were resolved approximately 6 cm into a pre-cast 4–12% Bis-Tris polyacrylamide gel (ThermoFisher Scientific). The lane was excised and cut in 8 approximately equal chunks and the proteins reduced, alkylated and digested in-gel. The resulting tryptic peptides were analysed by LC-MSMS using a Q Exactive coupled to an RSLCnano3000 (ThermoFisher Scientific). Raw files were processed using MaxQuant 1.5.2.8 using Andromeda to search a human Uniprot database (downloaded 03/12/14). Acetyl (protein N-terminus), oxidation (M) and deamidation (N/Q) were set as variable modifications and carbamidomethyl (C) as a fixed modification. SILAC data was loaded in R to process it with the Microarray-oriented limma package to call for differential expression [22], relying on the original normalisation processes produced by MaxQuant as reported previously [23, 24]. Three independent SILAC experiments using control U2OS and CHCHD4 (WT.cl1)-expressing cells involving parallel labelling were performed. Two independent SILAC experiments using shRNA control and CHCHD4 (shRNA) knockdown cells involving parallel labelling were performed.

### Sulforhodamine B (SRB) assay

Cells were plated in appropriate tissue culture vessels, and allowed to adhere overnight prior to treatment. Media was removed and cells were fixed with 10% trichloroacetic acid (TCA, #91228, Sigma-Aldrich) for 30 min. TCA was washed with water, wells were allowed to air dry, and then an excess of 0.4% (w/v) SRB (#S9012, Sigma-Aldrich) in 1% acetic acid (A/0400/PB15, ThermoFisher Scientific) was used to stain fixed cells for >10 min. Excess SRB was washed off with 1% acetic acid solution. Bound SRB was resuspended in a suitable volume of 10 mM Tris, and absorbance of solution measured at 570 nm. For proliferation assays, cells were plated on ‘day -1’ in triplicate in 12 well plates, and cultured in maintenance DMEM overnight, after which ‘day 0’ plates fixed with TCA. For galactose conditions, glucose media was replaced with glucose-free DMEM, supplemented with 4.5 g/L galactose, 10% FCS, penicillin, streptomycin and L-glutamine, and 1 mM pyruvate. Cells were then incubated for desired time points in either normoxia or hypoxia, followed by SRB assay. For drug sensitivity assays, cells were plated in triplicate columns in 96-well plates, and cultured in maintenance DMEM overnight. Appropriate wells were dosed with serial dilutions of compounds, including vehicle control wells. Cells were incubated for desired time points in either normoxia or hypoxia, followed by SRB assay.

### CI assay

For evaluating the direct inhibitory effect of compounds on CI activity from whole CI enzyme immunocaptured from purified bovine heart mitochondria the MitoTox™ Complex I OXPHOS Activity Microplate Assay (Abcam, ab109903) was used according to the manufacturer’s instructions. The EC_50_ for BAY 87-2243 in this assay was determined by calculating the dose at which 50% inhibition of CI activity was achieved. This was calculated as ~500 nM (see Fig. 4a).

### ATP assay

Cellular ATP concentrations were determined using CellTiter-Glo Luminescent Cell Viability Assay reagent and protocol, purchased from Promega (G7570). Briefly, cells were incubated in parallel in appropriate culture vessels under the conditions and for the duration of time described in the figure legends. Cells were lysed by the addition of a volume of CellTiter-Glo reagent equal to the volume of the culture media, and incubated for 2 min on an orbital shaker at room temperature. 100 μL of each sample was loaded into a 96 well luminometry plate, and the luminescence of each sample was measured using a Tecan Infinite M200 Pro luminescent plate reader. Parallel cell incubations were assayed for relative cell number using the SRB assay, and the luminescence measurements were normalised to the respective SRB measurements.

### Live cell imaging

Cells were plated in glass-based black-walled multi-welled microscopy plates, and allowed to adhere overnight. Following treatments (as described in figure legends), plates sealed with air-tight adhesive film, and imaged on a heated stage, using a DMI4000 B inverted microscope (Leica).

### Flow Cytometry

For TMRM staining: mitochondria were harvested according to protocol detailed here, and resuspended in DMEM supplemented with 1 mM L-pyruvate and 200 nM TMRM, and incubated for 30 minutes at 37°C and 5% CO_2_. After staining, mitochondria were washed twice in PBS, before resuspension in PBS, followed by flow cytometric analysis using a Fortessa flow cytometer (BD Biosciences). For MitoView staining: cells were harvested by trypsinisation, and centrifugation, and washed twice in warmed PBS. Cells resuspended in warmed PBS, or warmed PBS containing 200 nM MitoView (green). Fluorescence intensity was measured using a FACSCalibur flow cytometer (BD Biosciences). For 2-NBDG uptake assay: cells in suspension were incubated in warmed (37°C) PBS, or warmed PBS containing 200 μM 2-NBDG for 20 minutes. Cells were then washed three times in warmed PBS followed by flow cytometric analysis using a Fortessa flow cytometer (BD Biosciences).

### Quantification of microscopy images

For MitoSOX staining: images were processed using Cell Profiler Image analysis software, and fluorescence intensity was calculated per field of view. Mean fluorescence intensity, standard error and significance were calculated for each condition.

### Quantification of western blots (densitometry)

Western blot signal intensity was measured per lane using ImageJ (NIH) analysis software. Sample protein band intensities were normalised to the load control protein β-actin (for whole cell lysates) or PHB1 (for mitochondrial fractions). Relative band intensities were calculated relative to internal control sample.

### Statistical analysis

All statistical analyses carried out using Microsoft Excel (2016 edition). Groups compared using unpaired, two-tailed student’s t-test assuming equal variance, with significance set at *p*<0.05. All error bars represent standard deviation from the mean (SD) except where stated. Sample numbers (n values) and significance marks indicated in figure legends. Area under the curve calculated for each condition or cell line, and significance of difference between groups calculated using student’s t-test.

## RESULTS

### Impact of CHCHD4 on the mitochondrial proteome

CHCHD4 is responsible for the binding and import of mitochondrial proteins containing a twin CX_9_C or CX_3_C motif including subunits of respiratory chain complexes [8, 10]. Previously, we demonstrated that CHCHD4 is required for maintaining basal cellular oxygen consumption rate (OCR) [3] and intracellular oxygenation in tumour cells [25]. Recently, we found that loss of CHCHD4 in renal carcinoma cells leads to decreased expression of a range of respiratory chain subunits including NDUFS3 (CI), SDHA (complex II, CII), UQCRC2 (complex III, CIII), and COXIV (CIV) [26], indicating that CHCHD4 controls the expression of a broader range of respiratory chain subunits than had previously been considered. Thus, we hypothesised that CHCHD4 affects basal cellular OCR and intracellular oxygenation by altering the expression levels of respiratory chain subunits. To investigate the extent to which CHCHD4 affects respiratory chain subunit expression and other mitochondrial proteins, we performed stable isotope labelling with amino acid in cell culture (SILAC) using L-lysine-^13^C_6_, ^15^N_2_ (Lys-8) and L-arginine-^13^C_6_, ^15^N_4_ (Arg-10) parallel double labelling of shRNA control and CHCHD4 (shRNA) knockdown cells, as well as CHCHD4 (WT)-expressing cells and control cells (Fig. 1a). Mitochondria were isolated and mass spectrophotometry performed. Approximately 1050-1450 proteins were analysed from up to three independent parallel double labelling SILAC experiments (Fig. 1b-c). Our analysis of proteins isolated from mitochondria of respectively labelled cells confirmed a significant decrease in CHCHD4 protein in CHCHD4 (shRNA) knockdown compared to shRNA control cells (Fig. 1b) and a significant increase in CHCHD4 protein in CHCHD4 (WT)-expressing cells compared to control cells (Fig. 1c). We observed respective decreased and increased expression of several known CHCHD4 substrates containing a twin-CX_n_C motif including COX17, COA6, TIMM8, CHCHD2 and TIMM13 [8, 27] as well as other putative CHCHD4 substrates (Fig. 1d and Table 1), while expression of proteins such as the outer mitochondrial membrane protein sorting and assembly machinery component 50 homologue (SAMM50) did not change. Most interestingly, we observed significant and dynamic changes in a broad range of respiratory chain subunits involving CI, CII, CIII and CIV (Fig. 1e-g, Additional file 1a-b and Table 1). Notably, we observed more significant protein changes in response to loss of CHCHD4 (Fig 1b, Table 1) compared to CHCHD4 overexpression (Fig 1c, Table 1), which was likely related to the respective differences in significance for CHCHD4 expression in each of these conditions (Table 1). Importantly, we consistently observed that fold changes of identified proteins in response to loss or gain of CHCHD4 expression were decreased and increased respectively across multiple SILAC experiments (Fig 1 and Table 1).

**Table 1:**
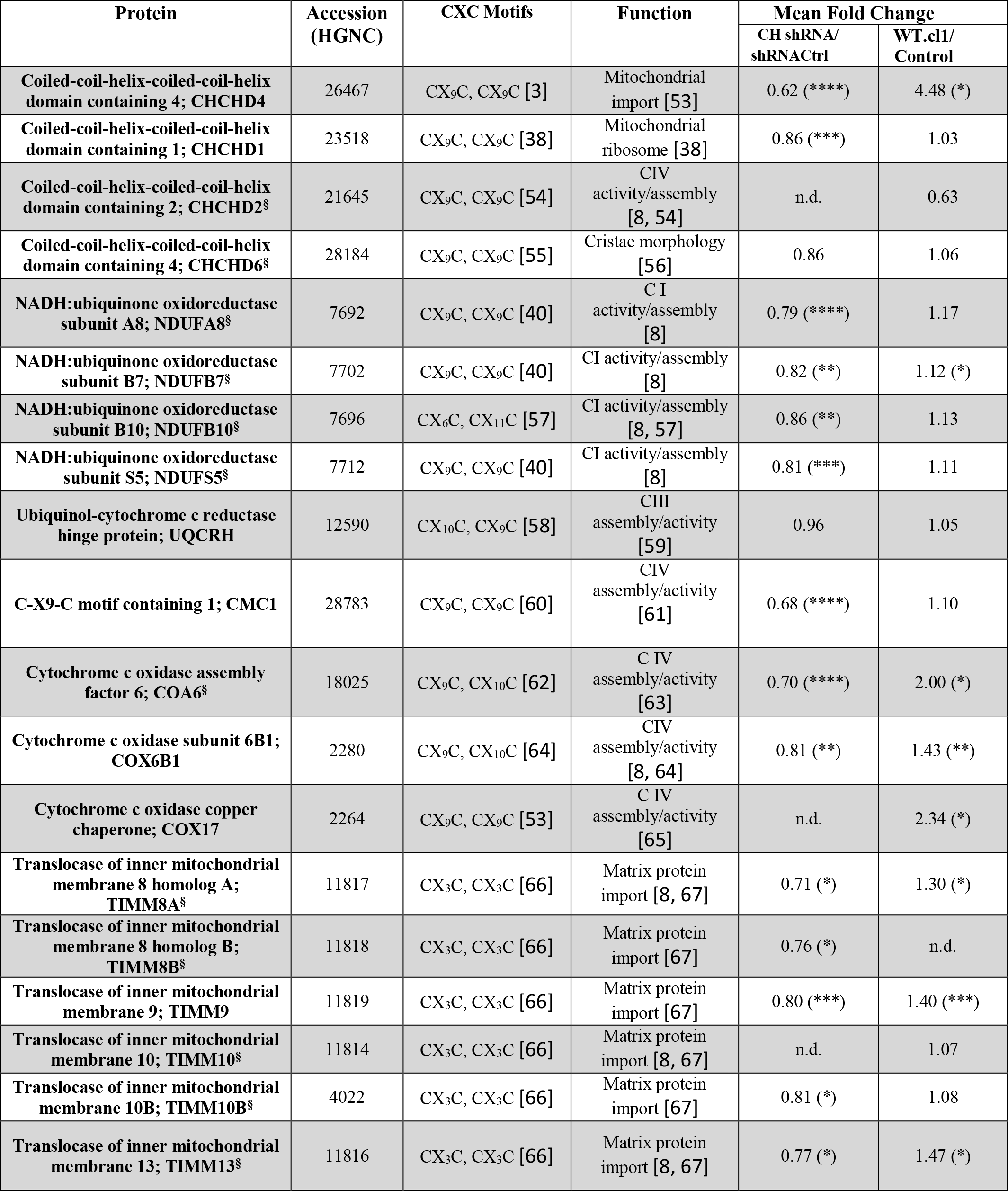
Twin CXnC motif-containing proteins identified in CHCHD4 (WT)-expressing cells and CHCHD4 (shRNA) knockdown cells using SILAC analysis. Table shows selected proteins, their respective twin CXnC motif and the fold change in their mitochondrial availability in CHCHD4 (shRNA) knockdown cells compared to control shRNA cells and CHCHD4 (WT)-expressing compared to control cells. Known CHCHD4 binding proteins are indicated (§). Representative mean fold changes in proteins observed in response to CHCHD4 knockdown were calculated from two parallel labelling analyses (CHCHD4 shRNA(H) vs shRNA Ctrl(L) and CHCHD4 shRNA(L) vs shRNA Ctrl(H)) from 2 independent SILAC experiments. Representative mean fold changes in proteins observed in response to elevated CHCHD4 expression were calculated from two parallel labelling analyses (WT(H) vs Ctrl(L) and WT(L) vs Ctrl(H)) from 3 independent SILAC experiments for all proteins shown, except CHCHD2 and TIMM8A where data was from 2 independent SILAC experiments. Proteins significantly changed are indicated * p<0.05, ** = p<0.01, *** = p< 0.001, **** =p< 0.0001. Proteins showing no reproducible change between experiments are indicated (n.d).

**Figure 1.**
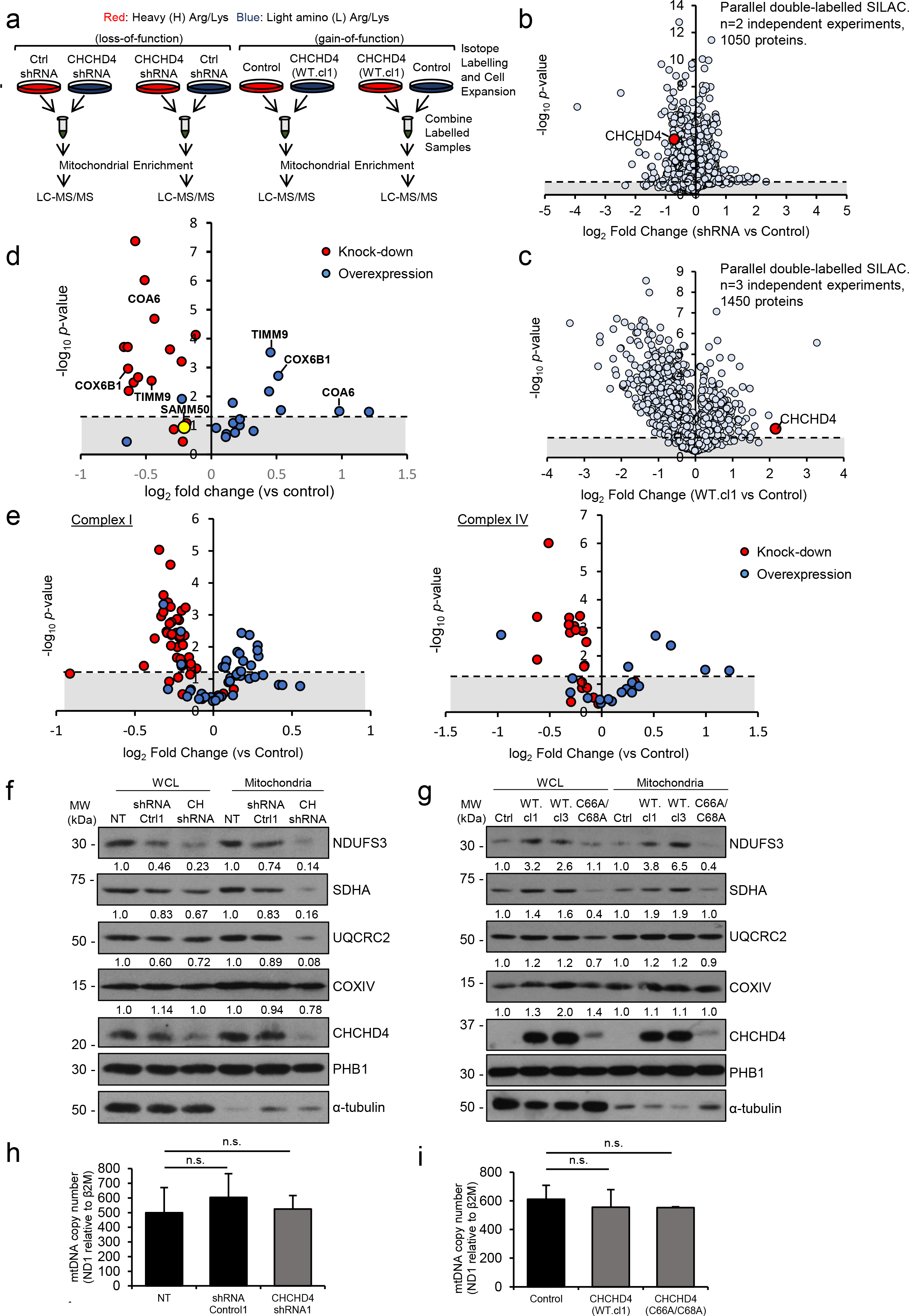
Impact of CHCHD4 on the mitochondrial proteome. **a** Schematic of SILAC procedure. HCT116 cells expressing control (non-targeting) shRNA or expressing CHCHD4-targeting shRNA and control U2OS or CHCHD4 (WT)-expressing U2OS cells (WT.cl1) cells were incubated in either L-lysine and L-arginine (light) or L-lysine-^13^C_6_, ^15^N_2_ (Lys-8) and L-arginine-^13^C_6_, ^15^N_4_ (Arg-10) (heavy) containing media for at least 5 cell divisions. Mitochondrial fractions were prepared and sample pairs were combined prior to LC-MS analysis. Data is representative of n=2 (b) and n=3 (c) independent SILAC experiments each involving parallel double labelling of cells. **b** Volcano plot showing log2 expression ratios of all detected proteins (1050) in enriched mitochondrial fractions from U2OS cells expressing CHCHD4-targeting shRNA compared to control (non-targeting) shRNA expressing cells. **c** Volcano plot showing log2 expression ratios of all detected proteins (1450) in enriched mitochondrial fractions from CHCHD4 (WT.cl1) expressing U2OS cells compared to control U2OS cells. CHCHD4 highlighted (red dot). **d** Volcano plot showing log2 expression ratios of known CHCHD4 substrates containing a twin-Cx_n_C motif identified from our SILAC analyses described in (a). SAMM50 is highlighted (yellow) as a mitochondrial protein that does not significantly change and is not a predicted CHCHD4 substrate. Dashed line denotes significance threshold, calculated by Student’s t-test, expressed as −log10 of calculated *p*-value. **e** Volcano plot, as in (b-c), showing changes in respiratory chain CI and CIV protein subunits from our SILAC analyses described in (a). Log2 expression ratios in WT.cl1 vs control U2OS (blue circles), and CHCHD4 shRNA1 vs control shRNA (red circles) shown. **f** Western blots show NDUFS3, SDHA, UQCRC2, COXIV and CHCHD4 protein levels in whole cell lysates (WCL) and mitochondrial fractions (Mitochondria) prepared from control HCT116 (NT) and HCT116 cells stably expressing an shRNA control vector (shRNA Ctrl1) or shRNA(1) targeting CHCHD4 (CH shRNA). PHB1 was used as a mitochondrial load control and α-tubulin was used as total load control. Densitometric ratio of each mitochondrial protein relative to PHB1 load is indicated. n=3. **g** Western blots show NDUFS3, SDHA, UQCRC2, COXIV and CHCHD4 protein levels in whole cell lysates (WCL) and mitochondrial fractions (Mitochondria) prepared from control U2OS (Ctrl), CHCHD4 (WT)-expressing (WT.cl1, WT.cl3) and CHCHD4 (C66A/C68A)-expressing cells. PHB1 was used as a mitochondrial load control and α-tubulin was used as total load control. Densitometric ratio of each mitochondrial protein relative to PHB1 load is indicated. n=3. **h-i** Graphs show mtDNA copy-number calculated as ratio of *mt-ND1* and *B2M* expression analysed by QPCR using total DNA isolated from cells described in (f) and (g) respectively. n=3; mean ± SD; n.s. = not significant.

To confirm our SILAC findings, we examined the expression of individual subunits of CI, CII, CIII and CIV in whole cell and enriched mitochondrial fractions by western analyses. Consistent with our SILAC analyses, we found reduced expression of individual subunits of CI (NDUFS3), CII (SDHA), CIII (UQCRC2) and CIV (COXIV) in enriched mitochondrial fractions isolated from stable CHCHD4 (shRNA) knockdown cells compared to shRNA control cells (Fig. 1f). Depletion of CHCHD4 using two independent siRNAs similarly reduced the expression of these respiratory chain subunits (Additional file 1b), as we have also shown in our recent study using renal carcinoma cells [28]. Furthermore, consistent with a previous report [29], we found that loss of CHCHD4 also reduced the expression of AIF protein, as well as GFER (Additional file 1c) without affecting their respective mRNA expression (Additional file 1d). Conversely, we found elevated levels of subunits of CI (NDUFS3), CII (SDHA), CIII (UQCRC2) and CIV (COXIV) in whole cell lysates and in enriched mitochondrial fractions isolated from CHCHD4 (WT)-expressing cells compared to control cells. However, these respiratory chain subunits were not increased in cells expressing a mutant form of CHCHD4 (C66A/C68A) in which the functionally critical cysteines of the substrate-binding CPC motif have been substituted by alanine, and is defective in mitochondrial localisation and import function [3, 15] (Fig. 1f). We observed no significant change in mitochondrial mass when CHCHD4 expression was manipulated as assessed either by measuring mitochondrial (mt)DNA copy-number (Fig. 1h-i) or by staining with the fluorescent mitochondrial marker, MitoView (Additional file 1e). Moreover, mitochondrial membrane potential was similar in mitochondria isolated from CHCHD4 (WT)-expressing cells, CHCHD4 (C66A/C68A)-expressing cells and control cells (Additional file 1f). Expression of the inner membrane protein prohibitin1 (PHB1) served as a control protein for these experiments as it did not change in response to either CHCHD4 (WT or C66A/C68A) expression or CHCHD4 knockdown (Fig. 1f-g). Collectively, our data indicate that CHCHD4 promotes global and dynamic effects on the expression of mitochondrial respiratory chain subunits.

### CHCHD4 promotes basal and adaptive metabolic responses, and provides a proliferative advantage to tumour cells

We have previously reported that overexpression of *CHCHD4* in human cancers correlates with hypoxia gene signature, increased tumour progression and metastasis, disease recurrence and poor patient survival [3]. As our SILAC analyses demonstrated significant changes in the expression of respiratory chain subunits, next we assessed the effect of CHCHD4 expression on tumour cell metabolism and growth rate in normoxia and hypoxia. In normoxia, we observed significantly decreased L-lactate levels (Fig. 2a) and extracellular acidification rate (ECAR) (Fig. 2b) in CHCHD4 (WT)-expressing cells compared with control or CHCHD4 (C66A/C68A)-expressing cells. Moreover, the rate of uptake of the fluorescent glucose analogue 2-NBDG (2-(N-(7-Nitrobenz-2-oxa-1,3-diazol-4-yl)Amino)-2-Deoxyglucose) was slower in CHCHD4 (WT)-expressing cells compared with control U2OS cells, while CHCHD4 (C66A/C68A)-expressing cells had higher uptake (Fig. 2c). Together, these data indicate that increased respiratory chain subunit expression correlates with decreased glycolysis in response to elevated CHCHD4 expression in normoxia, and decreased fermentation of pyruvate to lactate. In contrast to our findings in normoxia, we observed significantly increased L-lactate levels in CHCHD4 (WT)-expressing cells in hypoxia compared with control or CHCHD4 (C66A/C68A)-expressing cells (Additional file 2a), as we have shown previously [3]. Alongside this, we observed a corresponding change in the expression of the hypoxia inducible factor (HIF) target lactate dehydrogenase A (LDHA) (Additional file 2b), which converts pyruvate to lactate. Furthermore, we found that elevated CHCHD4 expression led to significantly increased tumour cell growth rate in both normoxia (Fig. 2d) and hypoxia (Additional file 2c), while the growth rate of CHCHD4 (C66A/C68A)-expressing cells was similar to control cells (Fig. 2d and Additional file 2c), indicating that increased expression of CHCHD4 in tumour cells provides a proliferative advantage in both normoxia and hypoxia. Taken together, our data suggest that CHCHD4-mediated effects on the respiratory chain regulate both basal (normoxic) and adaptive (hypoxic) metabolic tumour cell responses and growth rate.

**Figure 2.**
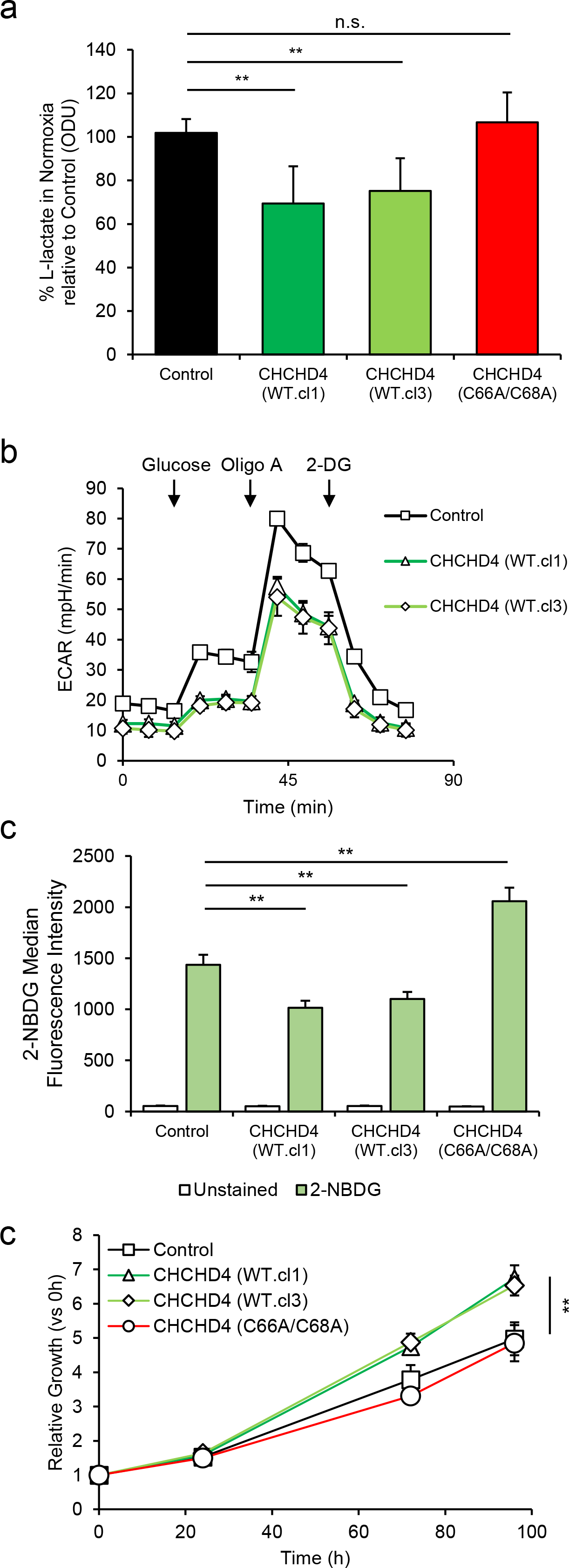
CHCHD4 promotes basal and adaptive metabolic responses, and provides a proliferative advantage to tumour cells. **a** Graph shows intracellular L-lactate levels from control U2OS cells, CHCHD4 (WT)-expressing cells (WT.cl1, WT.cl3) and CHCHD4 (C66A/C68A)-expressing cells, expressed as % of Control cells, measured using a cellular lactate assay. n=3; mean ± SD; n.s. = not significant, ** = *p*<0.01. **b** Graph shows ECAR (mpH/min) in cells described in (a), incubated in glucose-free basal media and measured using a Seahorse respirometer. Compounds were added as indicated to following final concentrations: glucose, 12.5 mM; oligomycin A, 1 μM; 2-deoxy-D-glucose, 50 mM. n=3; mean ± SD. **c** Chart shows intracellular fluorescence of cells described in (a) incubated with PBS (unstained) or 2-NBDG for 20 min. n=3; mean ± SD; ** = *p*<0.01. **d** Graph shows relative growth rate of cells described in (a), incubated over 96h. Total cell protein was assessed by SRB assay and used as a measure for cell growth. Relative growth was calculated for each time point relative to 0h. n=3; mean ± SD; ** = *p*<0.01.

### CHCHD4 confers increased tumour cell sensitivity to CI inhibitors

Homozygous deletion of *chchd4* in mice which is embryonic lethal, results in a decrease in whole CI expression [29], and human CI deficiency which is fatal, is associated with a cysteine mutation within the CI accessory subunit NDUFB10, which is a CHCHD4 substrate [30]. Our SILAC analyses of CHCHD4 (shRNA) knockdown and shRNA control cells showed a decrease in the expression of a range of CI accessory subunits (Fig. 1a, red dots). Conversely, CHCHD4 (WT)-expressing cells compared to control cells showed an increase in the expression of a range of CI subunits including NDUFS3, as well as NDUFA8, NDUFB10, and NDUFB7, which are considered to be CHCHD4 substrates [8, 27] (Fig. 1a, blue dots and Additional file 3). Given this relationship between CHCHD4 and CI biology, we hypothesised that elevated expression of CHCHD4 in tumour cells may render them more sensitive to respiratory chain OXPHOS inhibitors. Indeed, we found that CHCHD4 (WT)-expressing cells were significantly more sensitive to growth inhibition by the CI inhibitor rotenone compared to mutant CHCHD4 (C66A/C68A)-expressing or control cells in normoxia (Fig. 3a) and hypoxia (Additional file 4a), while there was no significant difference in the effects of sodium azide on growth across cell lines (Fig. 3b and Additional file 4b). As with sodium azide (Additional file 4c), rotenone also blocked HIF-1α protein (Additional file 4d). Together, our data show that the increased sensitivity of CHCHD4 (WT)-expressing cells to growth inhibition is specific to CI inhibition (rotenone) as opposed to CIV inhibition (sodium azide) and occurs irrespective of oxygen levels.

**Figure 3.**
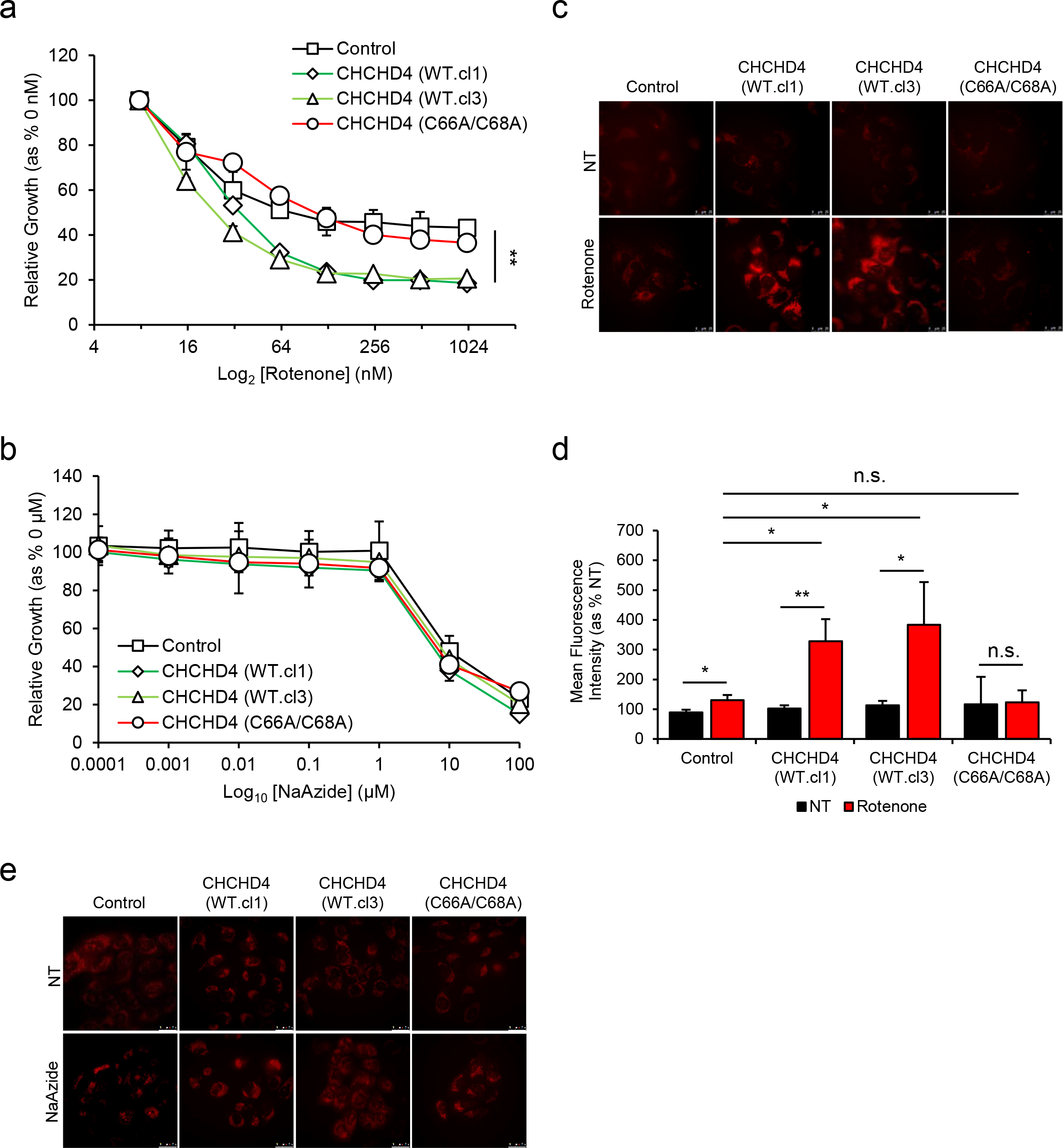
CHCHD4 confers increased tumour cell sensitivity to CI inhibitors. **a** Graph shows relative growth rate of control U2OS (control) cells, two independent CHCHD4 (WT)-expressing cell clones (WT.cl1, WT.cl3) and CHCHD4 (C66A/C68A)-expressing cells incubated for 72h in the absence or presence of rotenone using a 2-fold dilution series (top concentration, 1 μM). Total cell protein assessed by SRB assay was used as a measure of cell growth. Relative growth calculated for each time point for rotenone-treated relative to untreated. n=3; mean ± SD; * = *p*<0.05 (calculated from area under curve for Control vs CHCHD4 (WT.cl1) and (WT.cl3)). **b** As in (a), using a 10-fold dilution series of sodium azide (top concentration, 100 μM). n=3; mean ± SD. **c** Images of control U2OS cells, CHCHD4 (WT)-expressing cells (WT.cl1, WT.cl3) and CHCHD4 (C66A/C68A)-expressing cells, either untreated (NT) or treated with rotenone (500 nM) for 3h. MitoSOX Red ROS indicator (5 μM) was added for 30 min prior to live cell imaging by fluorescence microscopy. **d** Graph shows % mean fluorescence intensity quantified from images of cells described in (c). n=5 images per condition; mean ± SD; n.s. = not significant, * = *p*<0.05, ** = *p*<0.01. **e** Images of cells described in (c), either untreated (NT) or treated with sodium azide (NaAzide, 5 mM) for 3h. MitoSOX Red ROS indicator (5 μM) was added for 30 min prior to live cell imaging by fluorescence microscopy.

CI is a major source of mitochondrial ROS production, which can have profound effects on cellular viability through oxidative damage to lipids, proteins and nucleic acids [31]. Therefore next, we assessed mitochondrial ROS (superoxide) production using the ROS reporter MitoSOX. We found that basal ROS levels were similarly low in control and CHCHD4 (WT)-expressing cells (Fig. 3c-e). However, in response to rotenone, ROS levels were significantly increased in CHCHD4 (WT)-expressing cells compared to control or CHCHD4 (C66A/C68A)-expressing cells (Fig. 3c-d). Treatment of cells with sodium azide, did not lead to a change in mitochondrial ROS levels (Fig. 3e). Thus, increased sensitivity of CHCHD4 (WT)-expressing cells to CI inhibition (with rotenone) correlates with increased ROS production in these cells, suggesting that CHCHD4 determines tumour cell sensitivity to oxidative stress.

### CHCHD4 promotes mitochondrial ROS production in response to CI inhibition

To further explore the relationship between CHCHD4 and CI biology, we evaluated the effects of CI inhibition in CHCHD4 (WT)-expressing cells using the small molecule inhibitor BAY 87-2243 [32]. BAY 87-2243 was previously identified from a HIF-reporter screen and was shown to block HIF-1α by targeting CI without inhibiting CIII [32]. BAY 87-2243 exhibits potent antitumor activity in vivo [32–34], and has recently been shown to induce mitochondrial ROS through its CI inhibitory activity [35]. As expected, we found that BAY 87-2243 blocked whole CI enzyme activity at doses as low as 5 nM in vitro but was less potent than rotenone (Fig. 4a). We found that the EC_50_ dose (500 nM) of BAY 87-2243 for inhibiting whole CI enzyme activity (Fig. 4a) also blocked basal OCR by ~50% (Additional file 5a). Consistent with our findings with rotenone (Fig. 3a), we found that CHCHD4 (WT)-expressing cells were significantly more sensitive to growth inhibition by BAY 87-2243 treatment compared to control and mutant CHCHD4 (C66A/C68A)-expressing cells in normoxia (Fig. 4b) and hypoxia (Additional file 5b). Notably, the increased sensitivity of CHCHD4 (WT)-expressing cells to BAY 87-2243 was observed at very low nM doses (Fig. 4b and Additional file 5b) in normoxia and hypoxia. Moreover, there was no significant change in basal OCR (Additional file 5a) or total cellular ATP levels (Additional file 5c) at these very low nM doses of BAY 87-2243 treatment, suggesting that the effect of CI inhibition on growth inhibition is not mediated by changes at the level of intracellular oxygenation or bioenergetics. Indeed, similar to our findings with rotenone (Fig. 3c-d), ROS levels were significantly increased in CHCHD4 (WT)-expressing cells compared to control and CHCHD4 (C66A/C68A)-expressing cells in response to BAY 87-2243 treatment, even at a very low dose of 5 nM (Fig. 4c-d). The addition of a ROS scavenger, Trolox, significantly reduced ROS levels induced by BAY 87-2243 (Fig. 4c-d), and partially rescued the growth inhibitory effect of BAY 87-2243 treatment in CHCHD4 (WT)-expressing cells (Fig. 4e). Consistent with these data, we found that CHCHD4 (shRNA) knockdown cells were significantly less sensitive to BAY 87-2243 treatment (Fig. 4f), when cultured in glucose-free (galactose-containing) media which forces them to utilize the respiratory chain and produce ATP via OXPHOS [36]. Our data indicate that CHCHD4 confers tumour cell sensitivity to mitochondrial ROS produced by CI inhibition.

**Figure 4.**
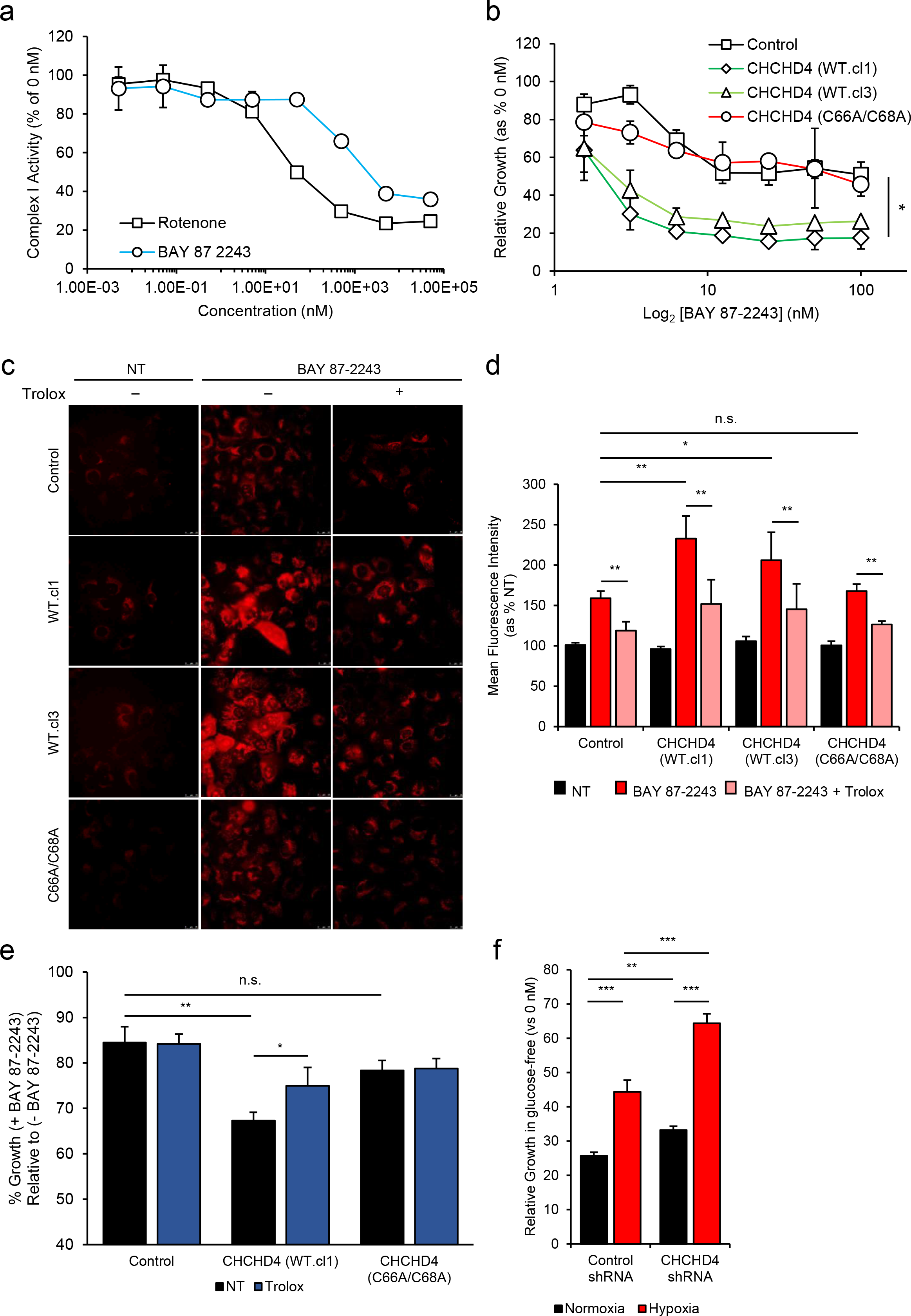
CHCHD4 promotes mitochondrial ROS production in response to CI inhibition. **a** Graph shows dose-response curve of whole CI activity in the presence of rotenone (black line) and BAY 87-2243 (blue line), using a 10-fold dilution series (top concentration 50 μM). **b** Graph shows relative growth rate of control U2OS cells, CHCHD4 (WT)-expressing cells (WT.cl1, WT.cl3) and CHCHD4 (C66A/C68A)-expressing cells incubated for 72h in the absence or presence of BAY 87-2243 using a 2-fold dilution series (top concentration, 100 nM). Total cell protein assessed by SRB assay was used as a measure of cell growth. Relative growth calculated for each time point for BAY 87-2243-treated relative to untreated. n=3; mean ± SD. * = *p*<0.05 (calculated from area under curve for Control vs CHCHD4 (WT.cl1) and (WT.cl3)). **c** Images of cells described in (b), either untreated (NT) or treated with of BAY 87-2243 (5 nM) for 3h, in the absence (-) or presence (+) of the ROS scavenger Trolox (100 μ MitoSOX Red ROS indicator (5 μ) was added for 30 min prior to live cell imaging by fluorescence microscopy. **d** Graph shows % mean fluorescence intensity quantified from images of cells described in (c), which were either untreated (NT) or treated with BAY 87-2243 (5 nM), or with BAY 87-2243 (5 nM) and Trolox (100 μM) (BAY + Trolox) for 3h. MitoSOX Red ROS indicator (5 μ imaging by fluorescence microscopy. n=5 images per condition; mean ± SD; n.s. = not significant, * = *p*<0.05, ** = *p*<0.01. **e** Graph shows % growth of cells treated without (−) or with (+) BAY 87-2243 (5 nM) for 24h, in the absence (NT, black bars) or presence of Trolox (100 μ) (blue bars). Total cell protein assessed by SRB assay was used as a measure of cell growth. Data represented as % growth of cells. n=3; mean ± SD. n.s. = not significant, * = *p*<0.05, ** = *p*<0.01. **f** Graphs show relative growth rates of control shRNA and CHCHD4 (shRNA1) cells in normoxia and hypoxia (1% O_2_) for 24h in the absence and presence of BAY 87-2243 (10 nM) incubated in glucose-free (galactose) containing DMEM. Relative absorbance for BAY 87-2243-treated cells was normalized against control (untreated, 0nM). n=2; mean ± SD; ** = *p*<0.01, *** = *p*<0.001.

### CHCHD4-mediated HIF-1α induction is blocked by NSC-134754 without affecting the respiratory chain

Previously, we have shown that CHCHD4 regulates HIF-1α protein induction and HIF signalling in hypoxia, and is required for tumour growth in vivo [3]. Here, we have found that elevated CHCHD4 expression increases tumour cell growth rate (Fig. 2c and Additional file 2c) and HIF-1α protein levels (Additional file 5c-d) in normoxia and hypoxia. Thus, next we assessed the contribution of HIF signalling to tumour cell growth rate observed in response to elevated CHCHD4 without targeting the respiratory chain. To do this, we used a small molecule HIF pathway inhibitor that we previously identified, named NSC-134754 [16]. Additional file 6a shows our revised chemical structure for NSC-134754 [20]. NSC-134754 blocks HIF activity in U2OS cells in normoxia and hypoxia within the sub-μM range (IC_50_ 0.25+0.05 μM). NSC-134754 mediates no direct inhibitory effects on basal OCR (Additional file 6b), does not inhibit whole CI enzyme activity (Fig. 5a), and does not induce mitochondrial ROS (Fig. 5b), collectively indicating that NSC-134754 blocks HIF signalling with no respiratory chain involvement. Moreover, in contrast to our growth inhibitor data observed with CI inhibitors (Fig. 3a and Fig. 4b), we found that NSC-134754 showed a comparable growth inhibitory profile in both control and CHCHD4 (WT)-expressing cells in normoxia and hypoxia (Fig. 5c-d). Furthermore, we found that elevated HIF-1α and HIF-2α protein induced in CHCHD4 (WT)-expressing in normoxia and hypoxia was blocked by NSC-134754 (Fig. 5f and Additional file 6c). Collectively our data show that our small molecule HIF signalling inhibitor, NSC-134754, can significantly block CHCHD4-mediated HIF-α protein induction without affecting the respiratory chain, highlighting the potential for targeting HIF signalling downstream of CHCHD4 and out-with the mitochondrial respiratory chain (Fig. 6).

**Figure 5.**
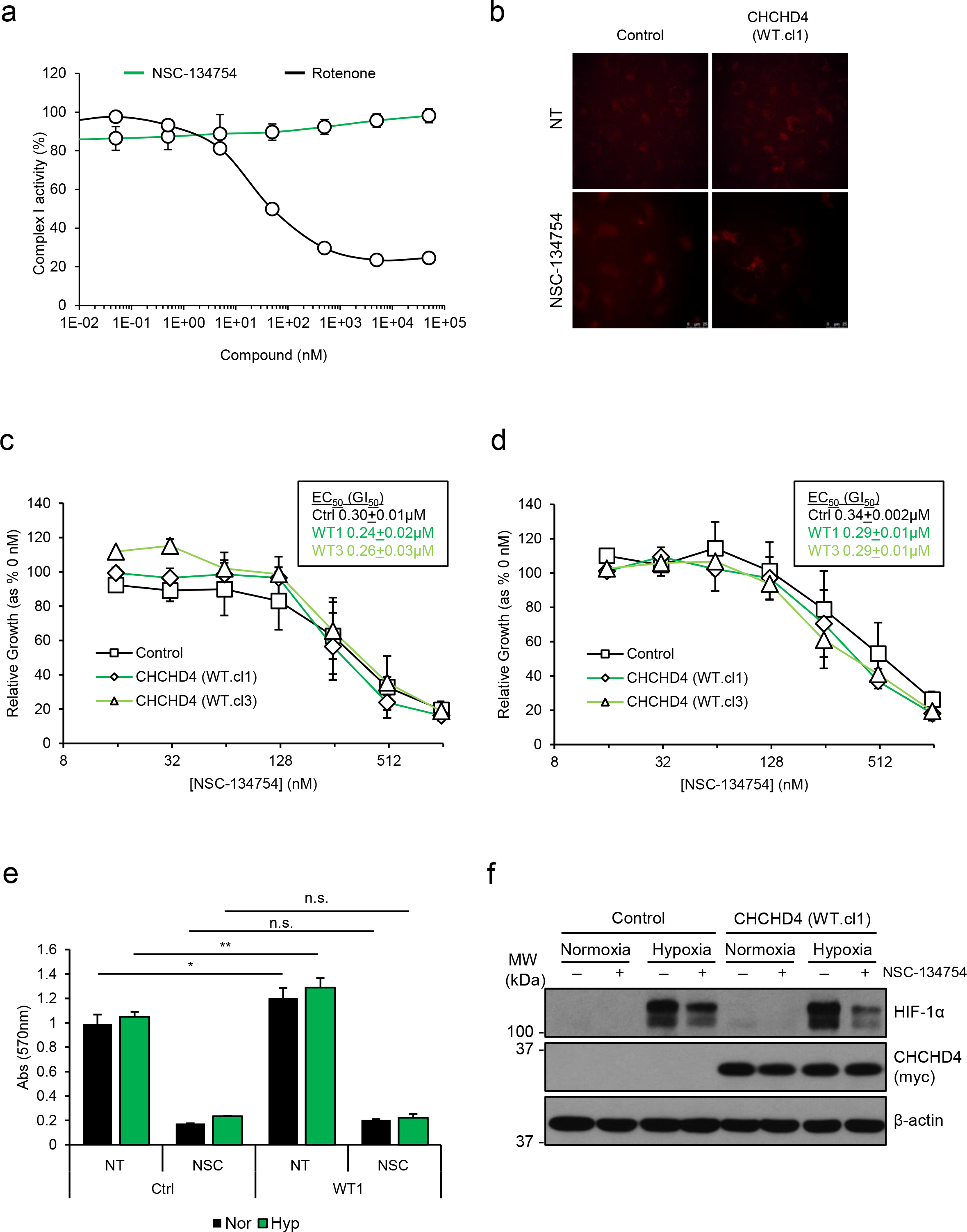
CHCHD4-mediated HIF-1α induction is blocked by NSC-134754 without affecting the respiratory chain. **a** Graph shows dose-response curve of whole CI activity in the presence of rotenone (black line) and NSC-134754 (green line), using a 10-fold dilution series (top concentration 50 μM). **b** Images of control U2OS cells, CHCHD4 (WT)-expressing cells (WT.cl1), either untreated (NT) or treated with NSC-134754 (1 μM) for 3h. MitoSOX Red ROS indicator (5 μM) was added for 30 min prior to live cell imaging by fluorescence microscopy. **c-d** Graphs show relative growth rate of control U2OS (control) cells, two independent CHCHD4 (WT)-expressing cell clones (WT.cl1, WT.cl3) and CHCHD4 (C66A/C68A)-expressing cells incubated for 72h in the absence or presence of NSC-134754 using a 2-fold dilution series (top concentration, 1 μM), in either normoxia (c), or hypoxia (1% O_2_) (d). Total cell protein assessed by SRB assay was used as a measure of cell growth. Relative growth calculated for each time point for NSC-134754-treated relative to untreated. EC_50_ values calculated using non-linear regression analysis. **e** Graph shows growth of control U2OS (control) cells and CHCHD4 (WT)-expressing cell clones (WT.cl1) for 72h in the absence (−) or presence (+) of NSC-134754 (1 μM), in normoxia (black bars) or hypoxia (1% O_2_, green bars). **f** Western blots show HIF-1α and exogenous CHCHD4 (myc) protein from control U2OS and CHCHD4 (WT)-expressing cells (WT.cl1) incubated in normoxia or hypoxia (1% O_2_) for 8h in the absence (−) or presence (+) of NSC-134754 (1 μM). β-actin was used as load control.

**Figure 6:**
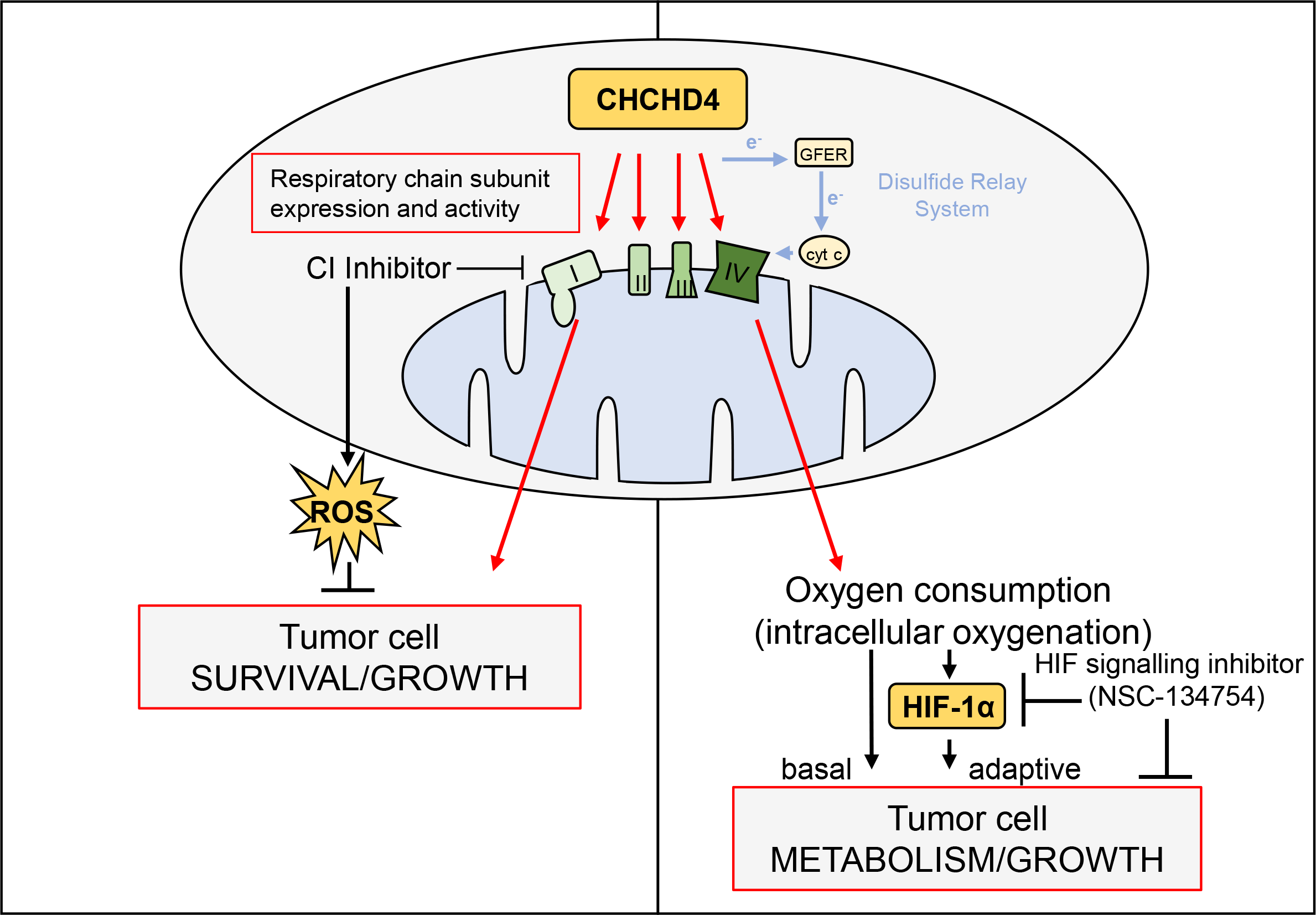
Schematic demonstrating proposed model of separable regulation of survival/proliferation and hypoxia responses by CHCHD4 through CI and CIV expression/activity respectively.

## DISCUSSION

Tumour cells rely on glycolysis and mitochondrial OXPHOS to survive [1]. Thus, mitochondrial metabolism has become an increasingly attractive area for investigation and therapeutic exploitation in cancer [2]. OXPHOS inhibitors are coming to the forefront for use in specific cancer types [37]. However, given the importance of mitochondrial OXPHOS for controlling intracellular oxygenation required for normal physiological processes, delineating whether or how OXPHOS inhibitors might provide a therapeutic window in cancer is crucially important.

Here we provide evidence that CHCHD4 expression in tumour cells is a critical determinant of the mitochondrial expression of a broad range of respiratory chain subunits including individual subunits of CI, CII, CIII and CIV, some of which are known CHCHD4 substrates (Table 1, Additional file 3 and [8]). These CHCHD4-regulated proteins contain both canonical (twin-CX_9_C or CX_3_C) and non-canonical (twin-CX_n_C) cysteine motifs, suggesting that CHCHD4 is capable of recognising and introducing disulfide bonds into proteins with a more diverse arrangement of cysteines than has previously been described [8]. In addition, we found changes in the mitochondrial expression of proteins previously identified as Mia40 substrates in yeast (e.g. CHCHD1 (Mrp10 in yeast) [38] (Table 1)).

Importantly, we found that increased expression of CHCHD4 or CHCHD4 knockdown increases or decreases respectively the expression of a number of supernumerary CI subunits involved in CI assembly (Table 1 and Additional file 3, [39]). Our data concur with a previous study showing that targeted deletion of *chchd4* in mice results in embryonic lethality by day E8.5, with null embryos displaying severe deficiency in total CI expression, accompanied by less severe defects in total CII-IV expression [29]. Interestingly, we identified at least four accessory CI subunits that are known CHCHD4 substrates (NDUFA8, NDUFB7, NDUFS5 and NDUFB10, Table 1). Three of these accessory CI subunits; NDUFA8, NDUFB7 [40] and NDUFS5 [41] have previously been identified as CHCHD4 substrates with canonical twin CX_9_C motifs [8], while a fourth subunit NDUFB10, contains a non-canonical cysteine motif (CX_6_C-CX_11_C) (Table 1), and has been shown recently to be mutated in a single case of fatal infantile lactic acidosis and cardiomyopathy in a way that renders it defective in binding to CHCHD4 [30]. A previous study describing the systematic genetic deletion of each of the 31 CI accessory subunits in cells shows that loss of either NDUFA8, NDUFB7, NDUFS5 or NDUFB10 has detrimental effects on the expression of other CI subunits, indicating that each of these CI subunits are strictly required for the assembly of whole CI [39]. Given the role of CHCHD4 import within the IMS, it is particularly interesting that these four accessory subunits of CI all reside within the IMS face of the CI structure [39, 42]. Therefore, it seems likely that CHCHD4 influences whole CI assembly and activity primarily through its ability to import multiple accessory CI subunits that are integrally involved in CI assembly [39]. Indeed, of the 30 nuclear-encoded CI subunits identified in our SILAC analyses (excluding those already confirmed as CHCHD4 substrates), 16 have two or more cysteine residues that could potentially be recognised and oxidised by CHCHD4 (Additional file 3). Intriguingly, we found that mitochondrial expression of the core CI subunit NDUFS3 [39] was also affected by modulation of CHCHD4 expression (Fig. 1 and Additional file 3). NDUFS3 knockdown has been shown to significantly decrease tumour cell proliferation [43] and overexpression of NDUFS3 is associated with breast cancer invasiveness [44]. NDUFS3 is not a known or putative CHCHD4 substrate (Additional file 3). However, reduced expression of NDUFS3 [45] and CI subunits as well as other respiratory chain complex defects have been observed upon loss or knockdown of apoptosis-inducing factor (AIF) [29, 46]. CHCHD4 has been shown to bind to AIF and is able to restore respiratory function in AIF-deficient cells [29]. Thus, CHCHD4 may also influence whole CI assembly and activity in tumour cells via other proteins involved in DRS function independently of its ability to import CI accessory subunits.

Collectively, our data highlight a fundamental relationship between CHCHD4 function and CI biology. Targeting CI as a therapeutic strategy in cancer has recently garnered considerable interest, due in part to the promising anticancer attributes of the antidiabetic drug metformin and the related biguanidine phenformin [1]. Metformin has been shown to block a variety of cellular and metabolic processes including CI activity [47, 48]. However, targeting respiratory chain components as a therapeutic strategy comes with considerable risk given the importance of the respiratory chain for normal cellular function. Tumour cells with elevated CHCHD4 expression and increased CI subunit expression and activity, produced significant levels of mitochondrial ROS (superoxide) upon treatment with CI inhibitors. However, elevated CHCHD4 expression did not render tumour cells more sensitive to growth inhibition by treatment with the CIV inhibitor sodium azide.

CI and CIII are considered the major sites for mitochondrial ROS (superoxide) production [49]. While we did not measure precisely where mitochondrial ROS (superoxide) was being produced upon CI inhibition in CHCHD4 (WT)-expressing cells, BAY 87-2243 has been shown to block CI without inhibiting CIII [32], and was recently shown to induce mitochondrial ROS through its CI inhibitory activity [35]. Interestingly, a recent study has implicated HIF-1 in the induction of the CI subunit NDUFA4L2 as mechanism to prevent overproduction of ROS during hypoxia [50]. However, we found no change in NDUFA4L2 expression in response to increased (or decreased) CHCHD4 expression in our SILAC analyses.

Metabolic adaptation to hypoxia in tumours is primarily mediated by HIFs and includes the diversion of glucose-derived carbons from the TCA cycle and the conversion of pyruvate to lactate, by upregulation of the enzymes pyruvate dehydrogenase kinase 1 (PDK1) and LDHA [51]. This reduces the reliance of cells on mitochondrial OXPHOS for the production of ATP. Indeed here and previously, we have shown that elevated expression of CHCHD4 in tumour cells increases hypoxic production of cellular lactate [3]. Paradoxically, here we found that elevated CHCHD4 expression in tumour cells leads to significantly reduced levels of cellular lactate in normoxia and no significant change in *PDK1* (data not shown), which is consistent with an increased respiratory drive in these cells. It is highly possible that HIF transcriptionally-dependent (negative) regulatory effects on mitochondrial function are negated in tumour cells expressing elevated CHCHD4 in normoxia due to its profound drive of mitochondrial function despite low levels of constitutive HIF-1α These observations indicate the possibility that CHCHD4 expression confers a double advantage to proliferating tumour cells, by stimulating more efficient glucose utilisation in both the presence and absence of oxygen. Consistent with this idea, we found that tumour cells with elevated CHCHD4 expression exhibit significantly increased proliferative capacity in normoxia and hypoxia, suggesting that CHCHD4 is capable of promoting tumour cell growth and metabolic adaptive responses, potentially through increased respiratory drive and HIF-mediated signalling respectively. Indeed, we found that our small molecule inhibitor of HIF signalling, NSC-134754 [16, 18-20] which exhibits anti-tumour activity in vivo [18, 52], could significantly block tumour cell growth and HIF-α (HIF-1α and HIF-2α) protein in tumour cells expressing elevated CHCHD4 without mitochondrial respiratory chain involvement.

## CONCLUSIONS

Mitochondria function as bioenergetic and biosynthetic factories to support cell survival and proliferation, and are important regulators of intracellular oxygenation and responses to hypoxia. Our present study demonstrates that the mitochondrial protein CHCHD4 controls the mitochondrial expression of a broad range of respiratory chain subunits. This reduces the glycolytic demands of cultured tumour cells and provides a proliferative advantage. However, our study also demonstrates that elevated CHCHD4 expression renders tumour cells more sensitive to growth inhibition by CI inhibitors, in part by increasing ROS production. Furthermore, while elevated CHCHD4 increases HIF-α protein induction in hypoxia, this is also sensitive to respiratory chain inhibition. Respiratory chain OXPHOS inhibitors may offer a potential therapeutic route for the treatment of tumours with increased proliferative drive, but this comes with significant risk of toxicity. Our study shows that our HIF pathway inhibitor NSC-134754 potently inhibits the growth of tumour cells with elevated CHCHD4 expression, without influencing mitochondrial function or ROS production. Further exploration of the mechanism of action of NSC-134754 will be particularly interesting.

## DECLARATIONS

### Ethics approval and consent to participate

This study did not involve human participation, personal data or use of human tissue.

### Consent for publication

Not applicable.

### Availability of data and materials

Requests can be made to the corresponding author relating to materials generated in this study.

### Authors’ contributions

LWT designed and performed experiments, analysed data and contributed to writing the manuscript. JS generated stable CHCHD4 (shRNA) HCT116 and U2OS knockdown cell lines, designed and performed SILAC experiments. CE designed and performed CI inhibitor experiments, whole CI assays, and analysed data. SH and RA assisted with SILAC analysis. AA and HAH contributed to the NSC-134754 inhibitor experiments. MA provided the concept for the study, designed experiments, analysed data, wrote the manuscript and acquired funding. All authors edited and reviewed the manuscript.

## Acknowledgements

Thanks to all members of the Ashcroft laboratory, especially Rachel Morgan for technical support. We thank Christian Frezza (MRC/Hutchison Cancer Unit), Patrick Maxwell (University of Cambridge, School of Clinical Medicine, UK) and members of the MRC Mitochondrial Biology Unit for their interest in this study.

## Authors’ information

Not applicable.

## ADDITIONAL FILES

Additional files 1-8 include five additional figures and three additional tables.

## REFERENCES

1. Weinberg, S.E. and N.S. Chandel, Targeting mitochondria metabolism for cancer therapy. Nat Chem Biol, 2015. 11(1): p. 9–15.

2. Galluzzi, L., et al., Metabolic targets for cancer therapy. Nat Rev Drug Discov, 2013. 12(11): p. 829–46.

3. Yang, J., et al., Human CHCHD4 mitochondrial proteins regulate cellular oxygen consumption rate and metabolism and provide a critical role in hypoxia signaling and tumor progression. J Clin Invest, 2012. 122(2): p. 600–11.

4. Thomas, L.W., et al., CHCHD4 Regulates Intracellular Oxygenation and Perinuclear Distribution of Mitochondria. Frontiers in Oncology, 2017. 7(71).

5. Erdogan, A.J. and J. Riemer, Mitochondrial disulfide relay and its substrates: mechanisms in health and disease. Cell Tissue Res, 2017. 367(1): p. 59–72.

6. Chatzi, A., P. Manganas, and K. Tokatlidis, Oxidative folding in the mitochondrial intermembrane space: A regulated process important for cell physiology and disease. Biochim Biophys Acta, 2016. 1863(6 Pt A): p. 1298–306.

7. Backes, S. and J.M. Herrmann, Protein Translocation into the Intermembrane Space and Matrix of Mitochondria: Mechanisms and Driving Forces. Front Mol Biosci, 2017. 4: p. 83.

8. Petrungaro, C., et al., The Ca(2+)-Dependent Release of the Mia40-Induced MICU1-MICU2 Dimer from MCU Regulates Mitochondrial Ca(2+) Uptake. Cell Metab, 2015. 22(4): p. 721–33.

9. Longen, S., et al., Systematic analysis of the twin cx(9)c protein family. J Mol Biol, 2009. 393(2): p. 356–68.

10. Modjtahedi, N., et al., Mitochondrial Proteins Containing Coiled-Coil-Helix-Coiled-Coil-Helix (CHCH) Domains in Health and Disease. Trends Biochem Sci, 2016. 41(3): p. 245–60.

11. Bihlmaier, K., et al., The disulfide relay system of mitochondria is connected to the respiratory chain. J Cell Biol, 2007. 179(3): p. 389–95.

12. Banci, L., et al., MIA40 is an oxidoreductase that catalyzes oxidative protein folding in mitochondria. Nat Struct Mol Biol, 2009. 16(2): p. 198–206.

13. Chacinska, A., et al., Mitochondrial biogenesis, switching the sorting pathway of the intermembrane space receptor Mia40. J Biol Chem, 2008. 283(44): p. 29723–9.

14. Hofmann, S., et al., Functional and mutational characterization of human MIA40 acting during import into the mitochondrial intermembrane space. J Mol Biol, 2005. 353(3): p. 517–28.

15. Fischer, M., et al., Protein import and oxidative folding in the mitochondrial intermembrane space of intact mammalian cells. Mol Biol Cell, 2013. 24(14): p. 2160–70.

16. Chau, N.M., et al., Identification of novel small molecule inhibitors of hypoxia-inducible factor-1 that differentially block hypoxia-inducible factor-1 activity and hypoxia-inducible factor-1alpha induction in response to hypoxic stress and growth factors. Cancer Res, 2005. 65(11): p. 4918–28.

17. Bunz, F., et al., Disruption of p^53^ in human cancer cells alters the responses to therapeutic agents. J Clin Invest, 1999. 104(3): p. 263–9.

18. Baker, L.C., et al., The HIF-pathway inhibitor NSC-134754 induces metabolic changes and anti-tumour activity while maintaining vascular function. Br J Cancer, 2012. 106(10): p. 1638–47.

19. Carroll, V.A. and M. Ashcroft, Role of hypoxia-inducible factor (HIF)-1alpha versus HIF-2alpha in the regulation of HIF target genes in response to hypoxia, insulin-like growth factor-I, or loss of von Hippel-Lindau function: implications for targeting the HIF pathway. Cancer Res, 2006. 66(12): p. 6264–70.

20. Hickin, J.A., et al., The synthesis and structure revision of NSC-134754. Chem Commun (Camb), 2014. 50(10): p. 1238–40.

21. Venegas, V. and M.C. Halberg, Measurement of mitochondrial DNA copy number. Methods Mol Biol, 2012. 837: p. 327–35.

22. Ritchie, M.E., et al., limma powers differential expression analyses for RNA-sequencing and microarray studies. Nucleic Acids Res, 2015. 43(7): p. e47.

23. Zhao, S., et al., The Application of SILAC Mouse in Human Body Fluid Proteomics Analysis Reveals Protein Patterns Associated with IgA Nephropathy. Evid Based Complement Alternat Med, 2013. 2013: p. 275390.

24. Howden, A.J., et al., QuaNCAT: quantitating proteome dynamics in primary cells. Nat Methods, 2013. 10(4): p. 343–6.

25. Thomas, L.W., et al., CHCHD4 Regulates Intracellular Oxygenation and Perinuclear Distribution of Mitochondria. Front Oncol, 2017. 7: p. 71.

26. Briston, T., et al., VHL-mediated regulation of CHCHD4 and mitochondrial function. Front Oncol, 2018. in press.

27. Modjtahedi, N. and G. Kroemer, CHCHD4 links AIF to the biogenesis of respiratory chain complex I. Mol Cell Oncol, 2016. 3(2): p. e1074332.

28. Briston, T., et al., VHL-Mediated Regulation of CHCHD4 and Mitochondrial Function. Front Oncol, 2018. 8: p. 388.

29. Hangen, E., et al., Interaction between AIF and CHCHD4 Regulates Respiratory Chain Biogenesis. Mol Cell, 2015. 58(6): p. 1001–14.

30. Friederich, M.W., et al., Mutations in the accessory subunit NDUFB10 result in isolated complex I deficiency and illustrate the critical role of intermembrane space import for complex I holoenzyme assembly. Hum Mol Genet, 2017. 26(4): p. 702–716.

31. Vinogradov, A.D. and V.G. Grivennikova, Oxidation of NADH and ROS production by respiratory complex I. Biochim Biophys Acta, 2016. 1857(7): p. 863–71.

32. Ellinghaus, P., et al., BAY 87-2243, a highly potent and selective inhibitor of hypoxia-induced gene activation has antitumor activities by inhibition of mitochondrial complex I. Cancer Med, 2013. 2(5): p. 611–24.

33. Helbig, L., et al., BAY 87-2243, a novel inhibitor of hypoxia-induced gene activation, improves local tumor control after fractionated irradiation in a schedule-dependent manner in head and neck human xenografts. Radiat Oncol, 2014. 9: p. 207.

34. Schockel, L., et al., Targeting mitochondrial complex I using BAY 87-2243 reduces melanoma tumor growth. Cancer Metab, 2015. 3: p. 11.

35. Basit, F., et al., Mitochondrial complex I inhibition triggers a mitophagy-dependent ROS increase leading to necroptosis and ferroptosis in melanoma cells. Cell Death Dis, 2017. 8(3): p. e2716.

36. Crabtree, H.G., The carbohydrate metabolism of certain pathological overgrowths. Biochem J, 1928. 22(5): p. 1289–98.

37. Molina, J.R., et al., An inhibitor of oxidative phosphorylation exploits cancer vulnerability. Nat Med, 2018.

38. Longen, S., et al., The disulfide relay of the intermembrane space oxidizes the ribosomal subunit mrp10 on its transit into the mitochondrial matrix. Dev Cell, 2014. 28(1): p. 30–42.

39. Stroud, D.A., et al., Accessory subunits are integral for assembly and function of human mitochondrial complex I. Nature, 2016. 538(7623): p. 123–126.

40. Szklarczyk, R., et al., NDUFB7 and NDUFA8 are located at the intermembrane surface of complex I. FEBS Lett, 2011. 585(5): p. 737–43.

41. Angerer, H., et al., A scaffold of accessory subunits links the peripheral arm and the distal proton-pumping module of mitochondrial complex I. Biochem J, 2011. 437(2): p. 279–88.

42. Zhu, J., K.R. Vinothkumar, and J. Hirst, Structure of mammalian respiratory complex I. Nature, 2016. 536(7616): p. 354–358.

43. He, X., et al., Suppression of mitochondrial complex I influences cell metastatic properties. PLoS One, 2013. 8(4): p. e61677.

44. Suhane, S., D. Berel, and V.K. Ramanujan, Biomarker signatures of mitochondrial NDUFS3 in invasive breast carcinoma. Biochem Biophys Res Commun, 2011. 412(4): p. 590–5.

45. Shen, S.M., et al., AIF inhibits tumor metastasis by protecting PTEN from oxidation. EMBO Rep, 2015. 16(11): p. 1563–80.

46. Meyer, K., et al., Loss of apoptosis-inducing factor critically affects MIA40 function. Cell Death Dis, 2015. 6: p. e1814.

47. Liu, X., et al., Metformin Targets Central Carbon Metabolism and Reveals Mitochondrial Requirements in Human Cancers. Cell Metab, 2016. 24(5): p. 728–739.

48. Wheaton, W.W., et al., Metformin inhibits mitochondrial complex I of cancer cells to reduce tumorigenesis. Elife, 2014. 3: p. e02242.

49. Sena, L.A. and N.S. Chandel, Physiological roles of mitochondrial reactive oxygen species. Mol Cell, 2012. 48(2): p. 158–67.

50. Tello, D., et al., Induction of the mitochondrial NDUFA4L2 protein by HIF-1alpha decreases oxygen consumption by inhibiting Complex I activity. Cell Metab, 2011. 14(6): p. 768–79.

51. Nakazawa, M.S., B. Keith, and M.C. Simon, Oxygen availability and metabolic adaptations. Nat Rev Cancer, 2016. 16(10): p. 663–73.

52. Kioi, M., et al., Inhibition of vasculogenesis, but not angiogenesis, prevents the recurrence of glioblastoma after irradiation in mice. J Clin Invest, 2010. 120(3): p. 694–705.

