## Supplemental Figure 1 for "CHCHD4 confers metabolic vulnerabilities to tumour cells through its control of the mitochondrial respiratory chain"

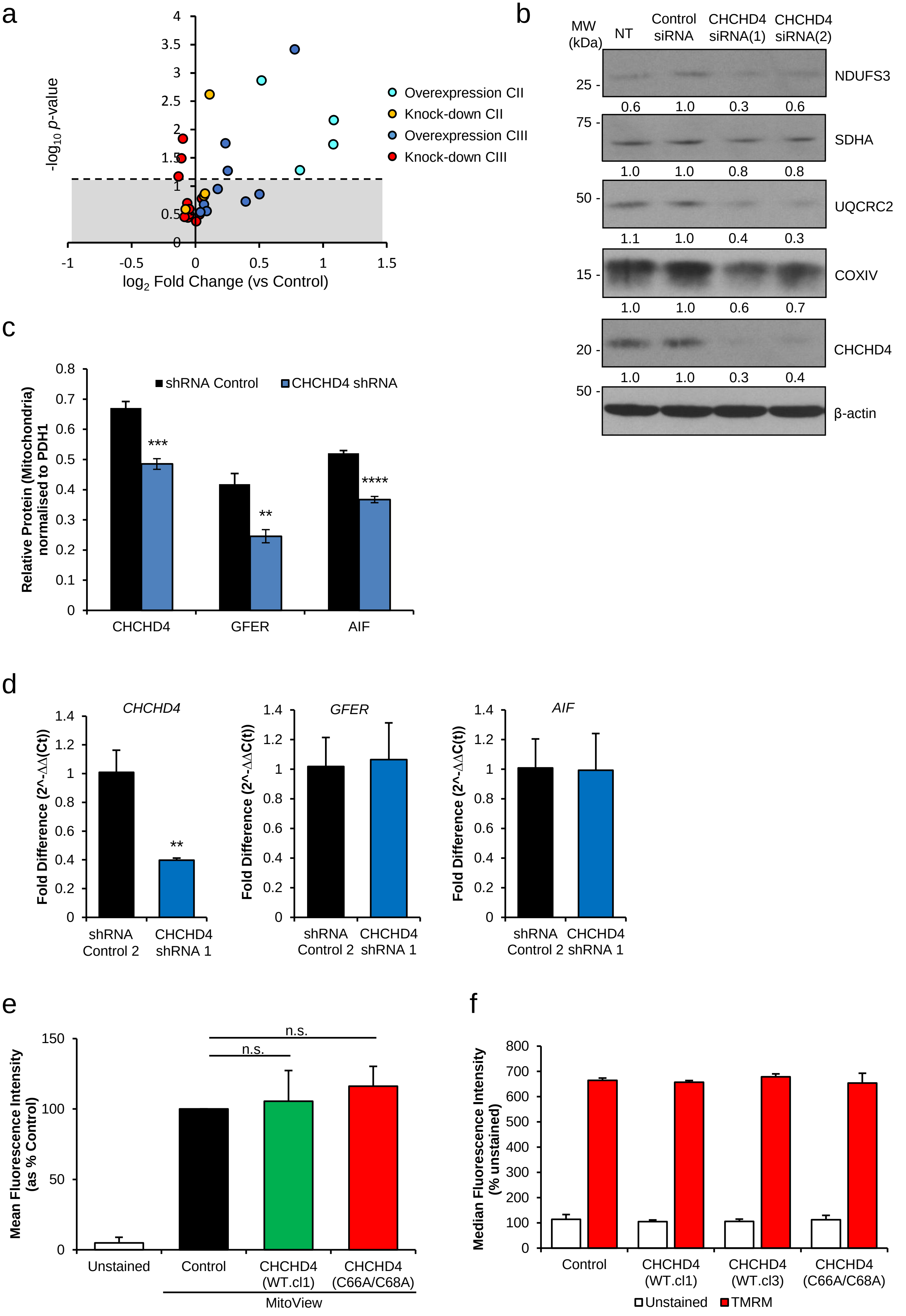


**Additional file 1.**

**Additional file 1.** Impact of CHCHD4 on the mitochondrial proteome. **a** Volcano plot shows relative expression of all detected subunits of complexes (C)II and III in CHCHD4 WT.cl1 expressing (overexpression) cells and CHCHD4 shRNA expressing (knock-down) cells, compared to control U2OS cells, detected by SILAC. **b** Western blots show NDUFS3, SDHA, UQCRC2, COXIV and CHCHD4 protein levels in control U2OS cells (NT) or transiently transfected with control siRNA, or two siRNAs targeting CHCHD4 (siRNA1 or siRNA2). β-actin used as load control. Densitometric ratio of each protein to β-actin is indicated. **c** Western blots show levels of AIF, GFER and CHCHD4 in subcellular fractions from control shRNA (shCtrl) expressing U2OS cells, and cells expressing CHCHD4-targeting shRNA (shCH). β-actin used as loading control. **c** Chart shows densitometric analysis of AIF, GFER and CHCHD4 protein levels relative to PHD1 protein from western blot analysis of subcellular fractions of control shRNA expressing U2OS cells (shRNA control), and cells expressing CHCHD4-targeting shRNA (CHCHD4 shRNA). n=3, mean ± SD **p<0.01, ***p<0.001, ****p< 0.0001. **d** Chart shows *CHCHD4*, *GFER*, and *AIFM1* (*AIF*) expression levels in cells described in (c). n=3, mean ± SD **p<0.01. **e** Graph shows mean fluorescence intensity measured by flow cytometry of control U2OS, CHCHD4 (WT)-expressing (WT.cl1, WT.cl3) and CHCHD4 (C66A/C68A)-expressing cells stained with 200 nM MitoView (green). mean ±SD; n=3; n.s. = not significant. **f** Graph shows median fluorescence intensity of unstained (white bars) and TMRM stained (red bars) mitochondria, isolated from control U2OS, CHCHD4 (WT)-expressing (WT.cl1, WT.cl3) and CHCHD4 (C66A/C68A)-expressing cells, prior to SILAC analyses described in Fig. 1a-e, western blots described in Fig. 1f-g. n=3; mean ± SD; n.s. = not significant.
