## Supplemental Figure 2 for "CHCHD4 confers metabolic vulnerabilities to tumour cells through its control of the mitochondrial respiratory chain"

a

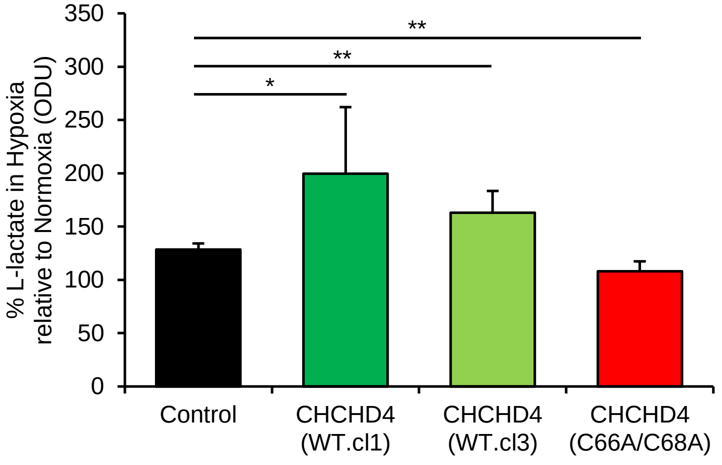


b

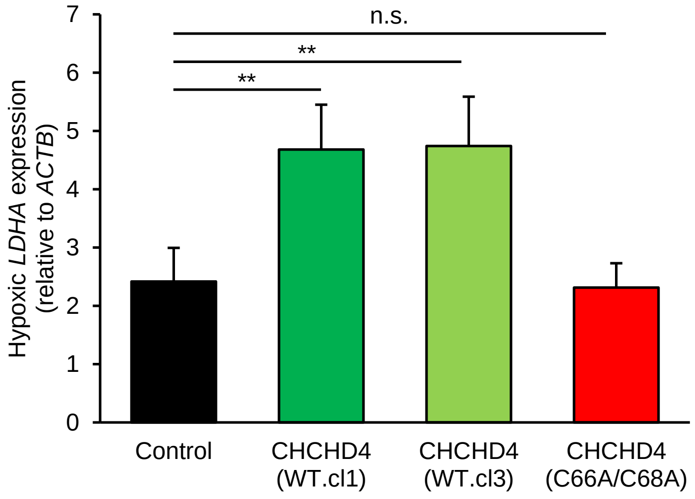


c

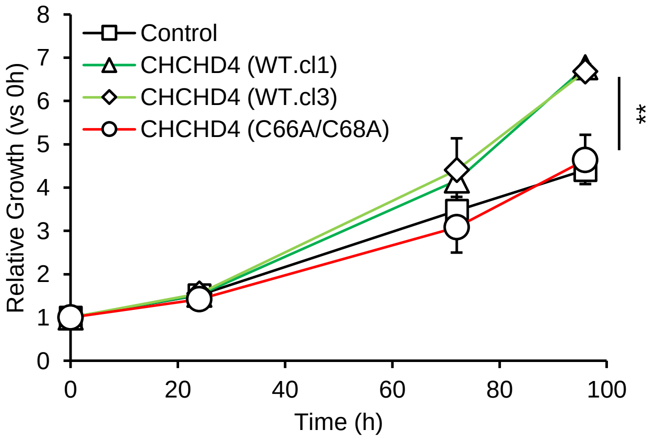


**Additional file 2.**

**Additional file 2.** CHCHD4 promotes basal and adaptive metabolic responses, and provides a proliferative advantage to tumour cells. **a** Graph shows total cellular L-lactate levels in control U2OS, CHCHD4 (WT)-expressing (WT.cl1, WT.cl3) and CHCHD4 (C66A/C68A)-expressing cells incubated in hypoxia (1% $O_2$) for 16h. Cellular L-lactate levels assayed spectrophometrically. Hypoxic L-lactate levels expressed as percentage of normoxic value. n=3; mean ± SD; * = $p<0.05$. ** = $p<0.01$. **b** Graph shows expression of *LDHA* analysed by QPCR using total RNA isolated from cells as in (a). n=3; mean ± SD; n.s. = not significant, ** = $p<0.01$. **c** Graph shows relative growth rate of cells described in (a), incubated over 96h in hypoxia (1% $O_2$). Total cell protein was assessed by SRB assay and used as a measure for cell growth. Relative growth was calculated for each time point relative to 0h. n=3; mean ± SD; ** = $p<0.01$.
