## Supplemental Figure 3 for "CHCHD4 confers metabolic vulnerabilities to tumour cells through its control of the mitochondrial respiratory chain"

| Subunit | Total Residues | Cysteine Residues | Mean Fold Change |
|---|---|---|---|
| NDUFA2 | 172 | 8 | 1.18 |
| NDUFA3 | 137 | 4 | 1.08 |
| NDUFA5 | 172 | 5 | 1.11 |
| NDUFA6 | 106 | 4 | 0.80 |
| NDUFA7 | 99 | 2 | 1.13 |
| NDUFA8* | 84 | 0 | 1.17 |
| NDUFA9 | 116 | 1 | 1.07 |
| NDUFA10 | 154 | 2 | 1.17 |
| NDUFA11 | 113 | 1 | 1.12 |
| NDUFA12 | 377 | 3 | 1.11 |
| NDUFA13 | 355 | 6 | 1.47 |
| NDUFAF2 | 228 | 8 | 0.87 |
| NDUFAF3 | 145 | 0 | n.d. |
| NDUFAF4 | 144 | 0 | 0.95 |
| NDUFB3 | 169 | 0 | 1.06 |
| NDUFB4 | 184 | 4 | 1.33 |
| NDUFB5 | 175 | 1 | 0.82 |
| NDUFB6 | 98 | 0 | 0.96 |
| NDUFB7* | 129 | 1 | 1.12 |
| NDUFB8 | 189 | 0 | 1.03 |
| NDUFB9 | 128 | 0 | 1.04 |
| NDUFB10* | 172 | 3 | 1.13 |
| NDUFB11 | 179 | 4 | 0.90 |
| NDUFC2 | 153 | 1 | 1.36 |
| NDUFS1 | 119 | 1 | 1.22 |
| NDUFS2 | 727 | 18 | 1.14 |
| NDUFS3 | 463 | 7 | 1.25 |
| NDUFS4 | 264 | 2 | 1.22 |
| NDUFS5* | 175 | 0 | 0.89 |
| NDUFS6 | 124 | 6 | 0.94 |
| NDUFS7 | 206 | 4 | 1.08 |
| NDUFS8 | 210 | 9 | 1.05 |
| NDUFV1 | 464 | 12 | 1.21 |
| NDUFV2 | 249 | 5 | 1.00 |

**Additional file 3: CHCHD4-mediated changes in the expression of nuclear encoded CI subunits identified using SILAC analysis.** Table shows mean fold change in the expression of CI subunits in response to elevated CHCHD4 expression. Data were calculated from two parallel labelling analyses (WT(H) vs Ctrl(L) and WT(L) vs Ctrl(H)) from 3 independent SILAC experiments. Total number of residues (amino acids) for proteins including the number of cysteine residues, and known CHCHD4 substrates (*) are indicated.
