## Supplemental Figure 4 for "CHCHD4 confers metabolic vulnerabilities to tumour cells through its control of the mitochondrial respiratory chain"

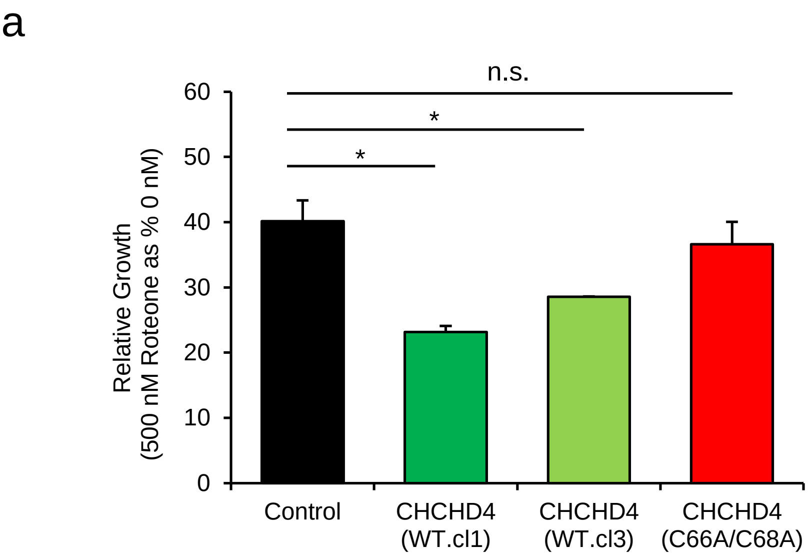


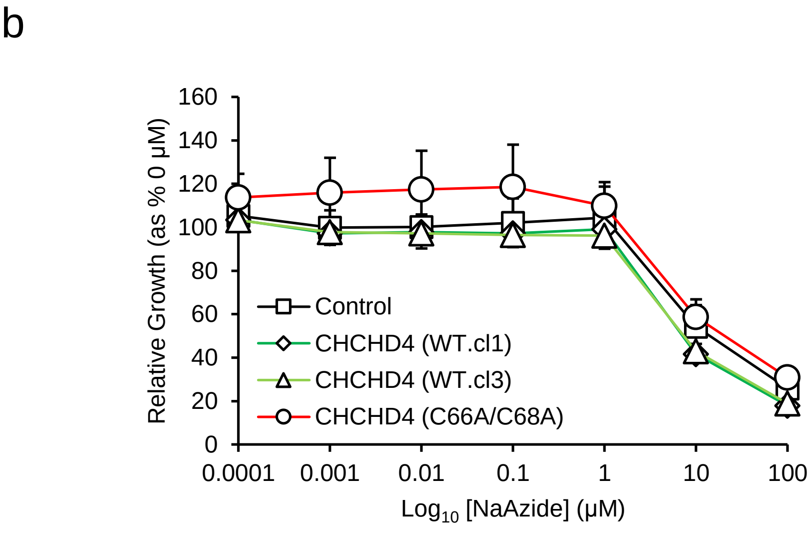


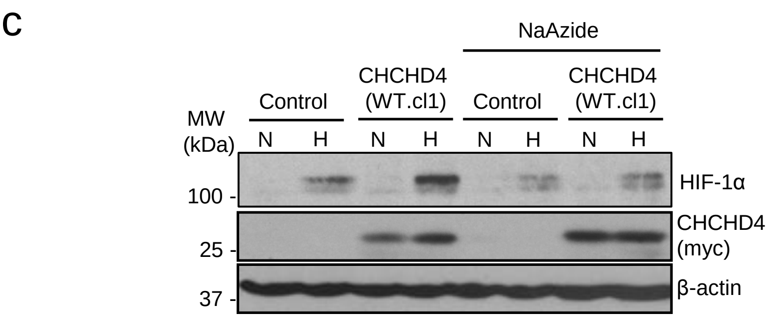


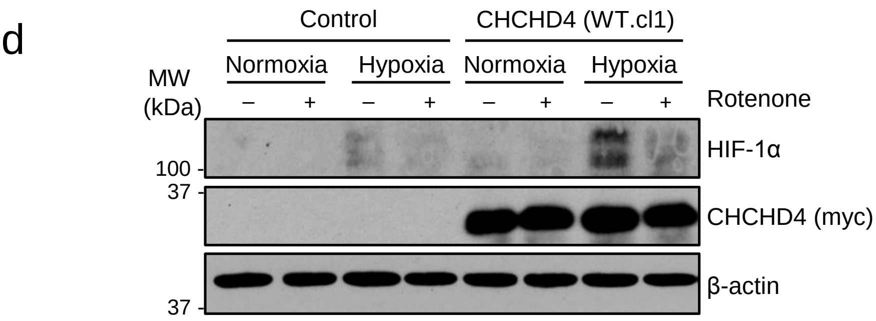


**Additional file 4.**

**Additional file 4.** CHCHD4 confers increased tumour cell sensitivity to CI inhibitors. **a** Graph shows relative growth rate of control U2OS (control) cells, two independent CHCHD4 (WT)-expressing cell clones (WT.cl1, WT.cl3) and CHCHD4 (C66A/C68A)-expressing cells incubated for 72h in hypoxia (1% $O_2$), in the absence or presence of rotenone (500 nM). Total cell protein assessed by SRB assay was used as a measure of cell growth. Relative growth calculated for rotenone-treated relative to untreated. n=3; mean ± SD; n.s. = not significant, * = $p<0.05$. **b** Graph shows relative growth rate of control U2OS (control) cells, two independent CHCHD4 (WT)-expressing cell clones (WT.cl1, WT.cl3) and CHCHD4 (C66A/C68A)-expressing cells incubated for 72h in hypoxia (1% $O_2$), in the absence or presence of sodium azide using a 10-fold dilution series (top concentration, 100 µM). Total cell protein assessed by SRB assay was used as a measure of cell growth. Relative growth calculated for rotenone-treated relative to untreated. n=3; mean ±SD. **c-d** Western blots show HIF-1α and CHCHD4 protein levels in control U2OS and CHCHD4 (WT)-expressing (WT.cl1) cells, untreated (-) or treated (+) with sodium azide (5 µM) (c) or rotenone (500 nM) (d) for 16h in normoxia or hypoxia (1% $O_2$). β-actin used as load control.
