## Supplemental Figure 5 for "CHCHD4 confers metabolic vulnerabilities to tumour cells through its control of the mitochondrial respiratory chain"

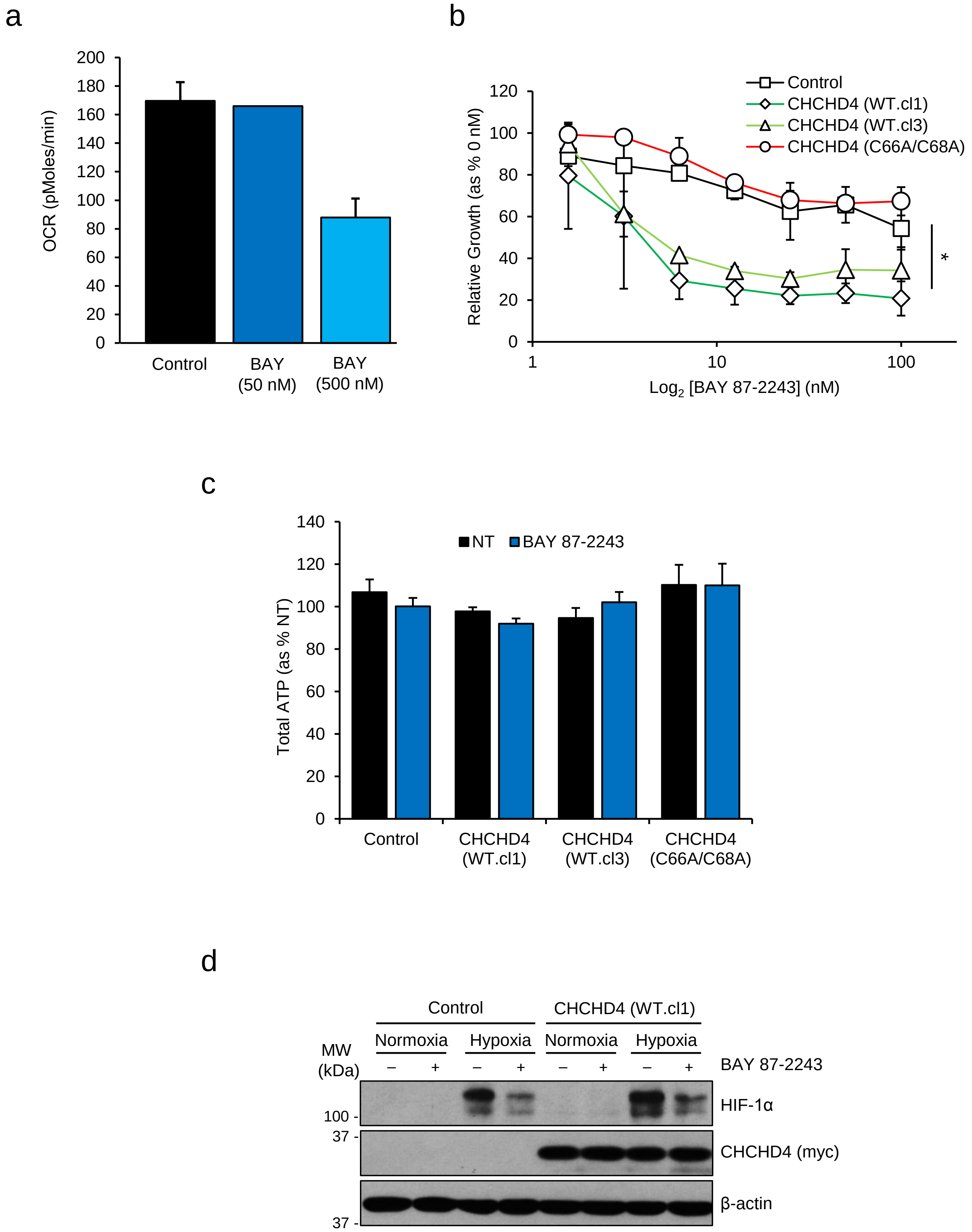


**Additional file 5.**

**Additional file 5.** CHCHD4 promotes mitochondrial ROS production in response to CI inhibition. **a** Graph shows basal OCR (pMoles/min) in control U2OS cells measured using a Seahorse respirometer in the absence (DMSO, control) or presence of BAY 87-2243 (BAY, 5 nM and 500 nM). n=3; mean ± SD. **b** Graph shows relative growth rate of control U2OS (control) cells, two independent CHCHD4 (WT)-expressing cell clones (WT.cl1, WT.cl3) and CHCHD4 (C66A/C68A)-expressing cells incubated for 72h in hypoxia (1% $O_2$), in the absence or presence of BAY 87-2243 using a 2-fold dilution series (top concentration, 100 nM). Total cell protein assessed by SRB assay was used as a measure of cell growth. Relative growth calculated for each time point for BAY 87-2243-treated relative to untreated (0 nM). n=3; mean ± SD. * = $p<0.05$ (calculated from area under curve for Control vs CHCHD4 (WT.cl1) and (WT.cl3)). **c** Graph shows total ATP (relative light units, RLU) in cells described in (b), treated with (white bars) or without (NT, black bars) BAY 87-2243 (5 nM) for 24h. RLU represented as % of untreated (NT) n=3; mean ± SD. **d** Western blots show HIF-1α and CHCHD4 protein levels in control U2OS and CHCHD4 (WT)-expressing (WT.cl1) cells, untreated (-) or treated (+) with BAY 87-2243 (1 µM) in normoxia or hypoxia (1% $O_2$) for 16h. β-actin used as load control.
