## Supplemental Figure 6 for "CHCHD4 confers metabolic vulnerabilities to tumour cells through its control of the mitochondrial respiratory chain"

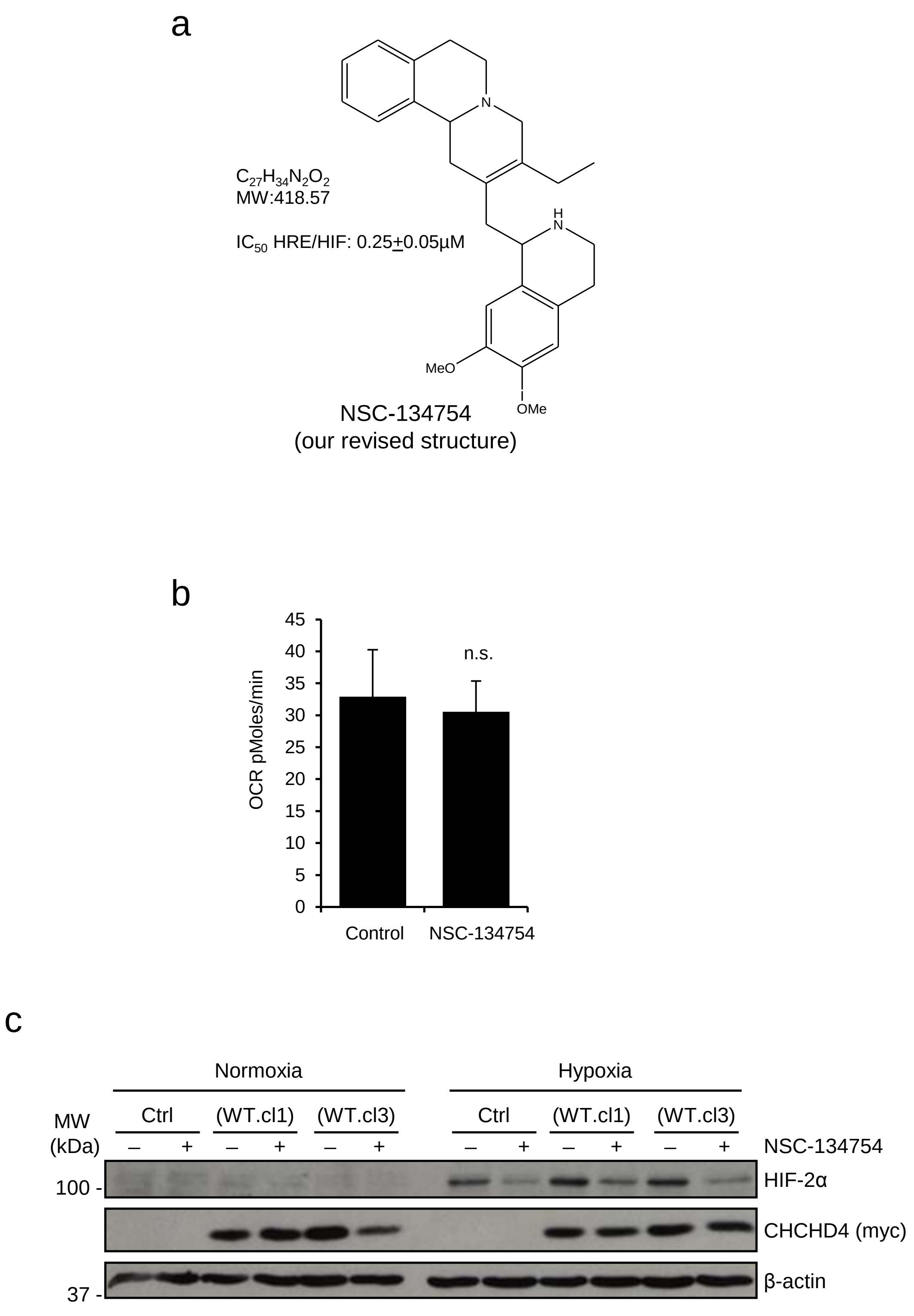


**Additional file 6.**

**Additional file 6.** CHCHD4-mediated HIF-α protein induction is blocked by NSC-134754 without affecting the respiratory chain. **a** Our revised chemical structure for NSC-134754. **b** Graph shows basal oxygen consumption rate (OCR, pMoles/min) in U2OS cells untreated or after treatment with NSC-134754 (10 μM) for 24h. n=3; mean ± SD; n.s. = not significant. **c** Western blots show HIF-2α and exogenous CHCHD4 (myc) protein levels in control U2OS and CHCHD4 (WT)-expressing cells (WT.cl1, WT.cl3), incubated in normoxia or hypoxia (1% $O_2$), untreated (NT) or treated with NSC-134754 (1 μM) for 8h. β-actin used as load control.
