## Supplemental Figure 7 for "CHCHD4 confers metabolic vulnerabilities to tumour cells through its control of the mitochondrial respiratory chain"

| ***siRNA/shRNA*** | | **Sequence** |
|---|---|---|
| ***CHCHD4* siRNA(1) target sequence** | | GAGGAAACGUUGUGAAUUA |
| ***CHCHD4* siRNA(2) target sequence** | | AAGAUUUGGACCCUUCCAUUC |
| ***CHCHD4* shRNA1 target sequence** | | UGUUCCUUGUUAUUCCGAA |
| ***CHCHD4* shRNA2 target sequence** | | GGAUCGAAUCAUAUUUGUA |

**Additional file 7: CHCHD4 siRNA and CHCHD4 shRNA sequences**. Table shows siRNA and shRNA sequences used for transient and stable knockdown of *CHCHD4* in cells.
