## Supplemental Figure 8 for "CHCHD4 confers metabolic vulnerabilities to tumour cells through its control of the mitochondrial respiratory chain"

| Gene | Forward Primer (5'-3') | Reverse Primer (5'-3') |
|---|---|---|
| ***CHCHD4*** | GAGCTGAGGAAGGGAAGGAT | AATCCATGCTCCTCGTATGG |
| ***GFER*** | GAGGAGTGTGCTGAAGACCT | CAGCTTGCGGTTCACTTCAT |
| ***AIF*** | GGCAAAATCGATAATTCTGTGGTTAGTC | CCACCAATTAGCAGGAAAGGAA |
| ***MT-ND1*** | GCCCCAACGTTGTAGGCCCC | AGCTAAGGTCGGGGCGGTGA |
| ***B2M*** | GAATGAGCGCCCGGTGTCCC | CCAAGCCAGCGACGCAGTG |

**Additional file 8: Q-PCR primer sequences.** Table shows independent forward and reverse primers used for the Q-PCR analysis of the genes indicated.
